# An intranasal, NLC-delivered self-amplifying RNA vaccine establishes protective immunity against pre-pandemic H5N1 and H7N9 influenza

**DOI:** 10.1101/2025.01.07.631792

**Authors:** Matthew R. Ykema, Michael A. Davis, Darshan N. Kasal, Madeleine F. Jennewein, Ethan Lo, Jasneet Singh, Samuel Beaver, Noah Cross, Eduard Melief, Sierra Reed, Christopher Press, Pauline Fusco, Julie Bakken, Corey Casper, Airn Tolnay Hartwig, Alana Gerhardt, Richard A. Bowen, Emily A. Voigt

**Affiliations:** Access to Advanced Health Institute (AAHI); Seattle, WA 98102, USA; Department of Medicine, University of Washington, Seattle, WA 98195, USA; Department of Global Health, University of Washington, Seattle, WA 98195, USA; Vaccine and Infectious Disease Division, Fred Hutch Cancer Center, Seattle, WA 98112, USA; Department of Biomedical Sciences, Colorado State University, Fort Collins, CO 80523, USA

## Abstract

Seasonal and pandemic influenzas are continuous threats to human health, requiring rapid development of vaccines to multiple evolving viral strains. New RNA vaccine technologies have the adaptability and manufacturability to facilitate pandemic preparedness but have limited flexibility in their route of administration, reducing the ability to establish local protective immune responses such as respiratory mucosal immunity. Here, we describe monovalent and bivalent self-amplifying RNA (saRNA) vaccines against A/Vietnam/1203/2004 H5N1 and A/Anhui/2013 H7N9. These saRNA vaccines express either H5 or H7 hemagglutinin and are formulated with a nanostructured lipid carrier (NLC) that permits both intramuscular (IM) and intranasal (IN) dosing. In mice, IM vaccination established systemic humoral and cellular responses but no detectable mucosal response, while IN administration induced robust systemic and mucosal immunity. The saRNA-NLC vaccines provided complete protection against morbidity and mortality in ferret challenge models, establishing this intranasally-administered saRNA-NLC vaccine platform as a potential pandemic response tool.

**ONE SENTENCE SUMMARY:** A self-amplifying RNA-NLC vaccine, delivered intranasally, induces robust mucosal immunity in mice and protects against H5N1 and H7N9 in ferrets

## INTRODUCTION

Influenza is a significant threat to human health, with seasonal circulation and pandemic outbreaks resulting in 9.3–41 million illnesses, 100,000–710,000 excess hospitalizations, and 4,900–51,000 deaths per year from 2010–2023 in the US alone (*1*). Of particular concern are zoonotic crossover events of “pre-pandemic” influenza strains that harbor the potential to gain human-to-human transmissibility (*2*). Avian influenza A strains have been detected in cows (H5N1), chickens (H5N2), and multiple avian species (H7N9), which increases the exposure risk to human hosts (*3, 4*). H5N1 influenza represents a particular concern given its current circulation in US bird and dairy cow populations (*3, 5*), its ability to infect multiple mammalian species (*6*), and its potential pathogenicity in humans resulting in high mortality in Africa, Asia, and the Middle East (*7*). As of January 2025, the CDC had confirmed 66 human H5N1 cases in the US across seven states, originating from cattle and poultry exposure sources (*8*). This risk is emphasized by an H5N1-associated death in the US in January 2025 (*9*). While not currently tracked as extensively as H5N1, H7N9 also presents a high potential risk due to its significant lethality in humans, up to 31.1%, from previous outbreaks (*10*). There are currently no major H7N9 outbreaks in animals, but multiple properties of this strain are of major concern, including the potential for animal-to-human transmission (*11*), transmission by respiratory droplets (*12*), and its ability to bind human sialic acid receptors (*13, 14*). Therefore, outbreaks of these variant strains pose a considerable risk for future pandemics.

Many vaccines have been developed to combat influenza (*15*), the majority of which are multivariant live-attenuated (LAIV) or inactivated (IIV) influenza vaccines against seasonal influenza strains that rely on traditional egg-based manufacturing methods. However, despite broad availability, vaccine effectiveness only ranges from 10 to 60% on a year-to-year basis and provides no protection against emerging pre-pandemic strains (*2*). Moreover, given the lead time required to isolate, culture, and formulate vaccines against emerging viral strains, LAIVs and IIVs are susceptible to antigenic drift and are severely limited in pandemic response (*16*).

RNA vaccine platforms overcome many of the limitations of traditional influenza vaccines by combining antigen flexibility with ease of manufacturing. As evidenced by the success of mRNA vaccines against SARS-CoV-2, which provided excellent protection against severe COVID-19, RNA-based vaccines exhibit properties necessary for pandemic preparedness and response, including establishment of robust immune responses. Recent work has also demonstrated their potential application in multivalent vaccine formulations (*17, 18*). However, there are major limitations with current RNA vaccines against respiratory pathogens, including their need for deep cold-chain storage (*19*), their inability to prevent mild infection in respiratory tissues (*20, 21*) possibly due to lack of established mucosal immunity (*22*), vaccine hesitancy (*23*), and rapid waning of vaccine-induced protective immunity (*24*).

Intranasally-administered vaccines are a promising tool for pandemic response. The ability to stimulate mucosal immunity at the site of natural respiratory viral infection may be key to improving vaccines’ ability to slow or halt a respiratory viral epidemic. In contrast to standard intramuscular (IM) vaccines, intranasal (IN) vaccines can stimulate mucosal immunity after local antigen exposure, with responses mediated by IgA antibodies and lung-resident memory T and B cells (*25, 26*), which can prevent early-stage infection and viral replication and shedding, thus minimizing chances for viral transmission (*27–29*). Intranasal vaccination devices also permit self-administration, the avoidance of needles, easier storage and transport (*30*), and simpler dosing of pediatric populations, thus increasing patient compliance and vaccine uptake. To date, the only FDA-approved IN vaccine is FluMist, a quadrivalent seasonal LAIV. Development of IN RNA vaccines would be a key tool for respiratory virus pandemic response, combining the mucosal immune stimulation of IN vaccines with the adaptability and production speed of RNA vaccines.

Here we report the application of a thermostable (*31, 32*) and intranasally-deliverable (*33*) self-amplifying RNA (saRNA)-nanostructured lipid carrier (NLC) vaccine platform towards the development of highly effective monovalent and bivalent saRNA-NLC vaccines against high-risk H5 and H7 influenza virus strains. We demonstrate the induction of both strong systemic and respiratory mucosal immune responses in mice that are unique to IN immunization. These vaccines also provide ferrets full protection from H5N1 and H7N9 viral-induced morbidity and mortality after high-dose challenge.

## RESULTS

### Intranasal administration of a monovalent H5 saRNA-NLC vaccine establishes dose-dependent systemic and mucosal immunity in mice

To establish proof-of-concept application of an saRNA-NLC vaccine platform against high-risk avian influenza, we generated a Venezuelan equine encephalitis virus (VEEV)-based saRNA expressing full-length H5 hemagglutinin (HA) derived from the prototypic A/Vietnam/1203/2004 H5N1 influenza strain. Vaccines were prepared by complexing H5-expressing saRNA with NLC, followed by biophysical characterization of the resulting vaccine complex size, RNA dose, and RNA integrity (**Supplementary Figure S1A-B**). This saRNA-NLC vaccine formulation was previously optimized to allow both IM and IN delivery.

We first verified the immunogenicity of this monovalent H5 saRNA-NLC vaccine in mice when administered IN by droplets using a pipette and/or IM via injection. C57BL/6 mice were prime/boost vaccinated 21 days apart either IN/IN (1, 5, or 10 µg), IM/IM (5 µg), or IM-prime, IN-boost (IM/IN, 5 µg) to examine the effects of dose and administration route on immunogenicity (**Figure 1A**). An alum-adjuvanted, inactivated whole virion influenza A/Vietnam/1203/2004 H5N1 vaccine (*34*) (“Baxter,” BEI Resources #NR-12143) delivering 2.5 µg of HA protein was dosed IM/IM as a positive vaccine comparator, while 5 µg of NLC-delivered saRNA expressing the non-immunogenic reporter protein secreted embryonic alkaline phosphatase (SEAP) dosed IN/IN served as a negative vector control. Reactogenicity was assessed by monitoring animal weights for 4 days post-vaccination (**Supplementary Figure S1C-D**).

**Figure 1.**
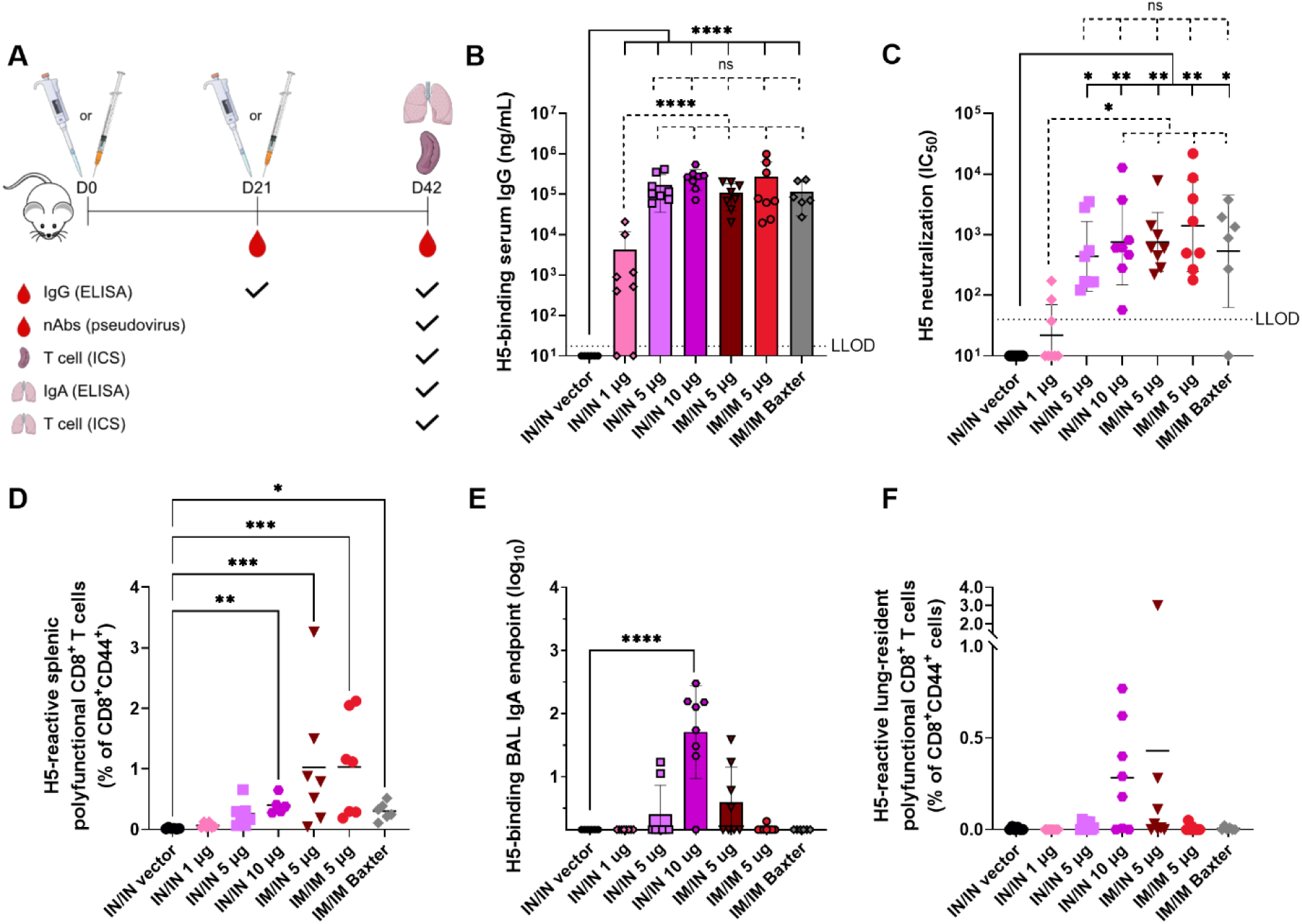
Intranasal administration of monovalent H5 saRNA-NLC establishes systemic and mucosal immune responses in mice. (A) Study design. Post-boost (B) serum H5-binding IgG titers, (C) serum H5 pseudovirus neutralization capacity (IC_50_), (D) splenic H5-reactive polyfunctional (IFN*γ*^+^ IL-2^+^ TNF*α*^+^) CD8^+^ T cells, (E) H5-binding IgA titers in BAL samples, and (F) lung-resident (CD69^+^ CD103^+^) H5-reactive polyfunctional (IFN*γ*^+^ IL-2^+^ TNF*α*^+^) CD8^+^ T cells. *n* = 6 for vector control group and *n* = 8 for experimental groups. All groups were sex balanced. Two statistical hypotheses were tested within each figure, with filled lines showing comparisons to the vector control and dotted lines representing tests between key experimental groups. (B, E) Statistical analyses performed on log-transformed data using one-way ANOVA with Šídák’s multiple comparison test. (C) Statistical analysis performed on log-transformed data using Kruskal-Wallis test with Dunn’s multiple comparisons. (D, F) Statistical analysis represents Kruskal-Wallis test with Dunn’s multiple comparisons. LLOD = lower limit of detection; ns = not significant. * *p* < 0.05, ** *p* < 0.01, *** *p* < 0.001, **** *p* < 0.0001. Associated post-prime IgG and post-boost CD4^+^ and CD8^+^ T cell data in **Supplementary Figure S2**.

All groups receiving at least 5 µg of H5 saRNA-NLC developed equivalent serum H5-binding IgG and neutralizing antibody (nAb) titers, despite differences in administration route, indicating that IN vaccination with saRNA-NLC establishes a robust systemic immune response comparable to IM or heterologous IM/IN immunization (**Figure 1B-C, post-boost; Supplementary Figure S2A, post-prime**). Additionally, regardless of administration route, 5 µg doses of H5 saRNA-NLC induced responses comparable to those seen in the Baxter vaccine control group. Proportions of splenic H5-reactive activated (CD44^+^) CD4^+^ T cells and CD8^+^ T cells similarly increased in a dose-dependent manner following IN vaccination (**Figure 1D, Supplementary Figure S2B-C**). Comparable frequencies of polyfunctional CD4^+^ and CD8^+^ T cells were observed for IN/IN 10 µg, IM/IN 5 µg, or IM/IM 5 µg H5 saRNA-NLC groups (p > 0.05), whereas the Baxter control H5N1 vaccine induced a negligible polyfunctional T cell response in the spleen. Notably, the heterologous IM/IN 5 µg regimen established splenic CD8^+^ T cell responses comparable to those induced by a double dose (10 µg) delivered IN (*p* > 0.05), indicating a potential advantage of a heterologous IM/IN prime/boost strategy. These data demonstrate that our H5 saRNA-NLC vaccine, delivered either IN or IM, stimulates systemic humoral and cellular immune responses in mice at least comparable to the adjuvanted Baxter vaccine.

In addition to overcoming vaccine hesitancy due to fear of needles and the potential for self-administration, an IN vaccine may also be advantageous for inducing localized respiratory mucosal immunity at the site of infection. Strikingly, development of both H5-binding IgA in bronchoalveolar lavage (BAL) samples and lung-resident (CD69^+^ CD103^+^) polyfunctional CD8^+^ T cells was observed in animals receiving at least one IN vaccination (**Figure 1E-F**). Little to no mucosal response was detected in either saRNA-NLC or Baxter IM-only vaccinated animals. Lung-resident (CD69^+^) polyfunctional CD4^+^ T cells and IFNγ^+^ CD8^+^ T cells followed a similar pattern (**Supplementary Figure S2D-E**). These data demonstrate that homologous IN/IN and heterologous IM/IN administrations of our H5 saRNA-NLC vaccine can establish both systemic and mucosal immune responses and that IN administration is necessary for the development of robust mucosal immunity.

### Intranasal administration of a monovalent H7 saRNA-NLC vaccine establishes dose-dependent systemic and mucosal immunity in mice

We next evaluated immunogenicity of a monovalent H7 saRNA-NLC candidate. Similar to our H5 saRNA-NLC vaccine, full-length H7 HA derived from A/Anhui/2013 H7N9 influenza was engineered into a VEEV-based saRNA, complexed with NLC, and characterized (**Supplementary Figure S3A-B**). Mice were prime/boost immunized with H7 saRNA-NLC via homologous (IN/IN or IM/IM) or heterologous (IM/IN) routes (**Figure 2A**), and reactogenicity was assessed as above (**Supplementary Figure S3C-D**). Unlike H5N1, no H7N9 vaccine comparator was readily available for this study.

**Figure 2.**
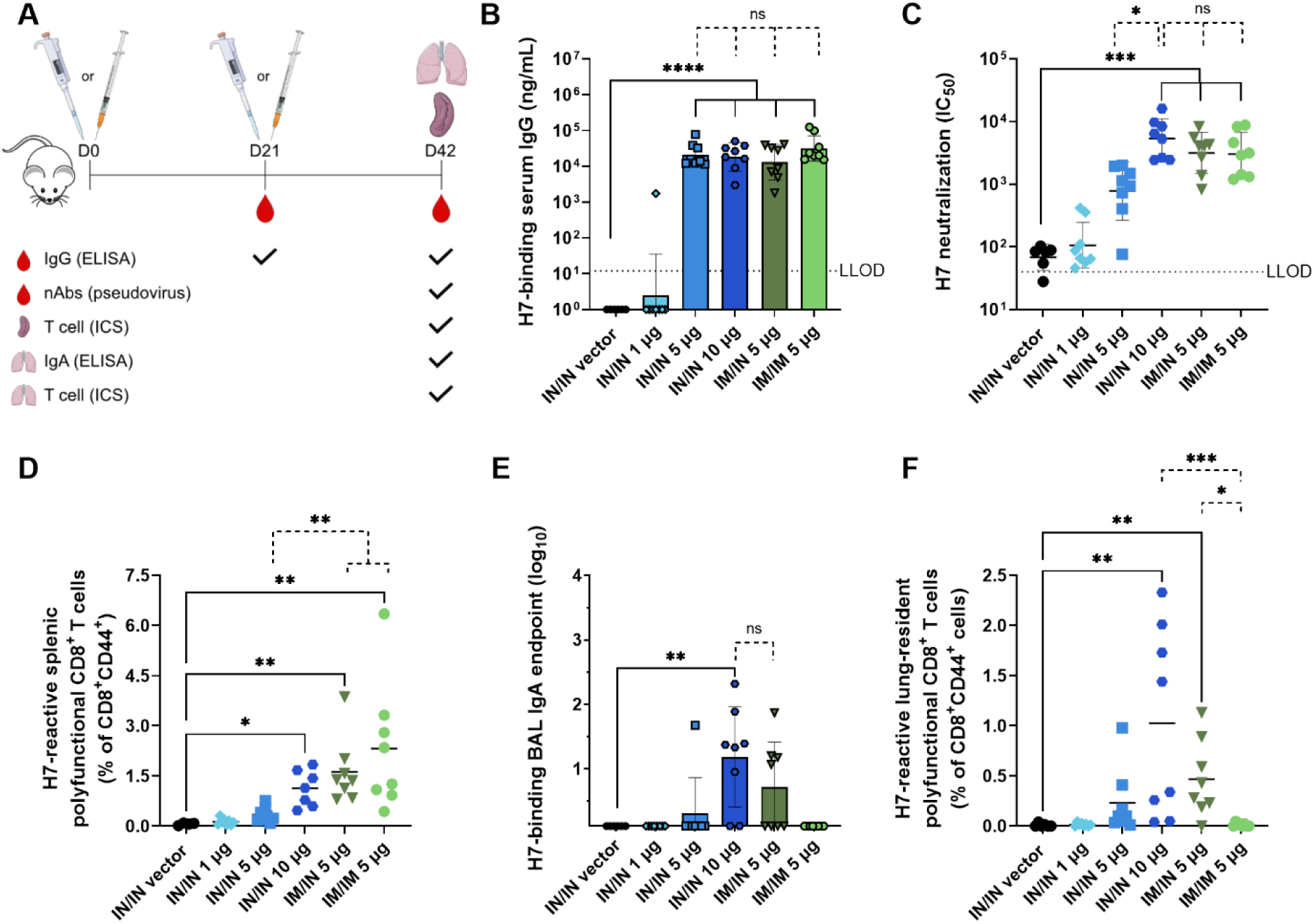
Intranasal administration of monovalent H7 saRNA-NLC establishes dose-dependent systemic and mucosal immune responses in mice. (A) Study design. Post-boost (B) serum H7-binding IgG titers, (C) serum H7 pseudovirus neutralization capacity (IC_50_), (D) splenic H7-reactive polyfunctional (IFN*γ*^+^ IL-2^+^ TNF*α*^+^) CD8^+^ T cells, (E) H7-binding IgA antibody titers in BAL samples, and (F) lung-resident (CD69^+^ CD103^+^) H7-reactive polyfunctional (IFN*γ*^+^ IL-2^+^ TNF*α*^+^) CD8^+^ T cells. For all assays, *n* = 6 animals for vector control group and *n* = 8 for experimental groups. All groups were sex balanced. Two statistical hypotheses were tested within each figure, with filled lines showing comparisons to the vector control and dotted lines representing tests between key experimental groups. (B, E) Statistical analyses performed on log-transformed data using one-way ANOVA with Šídák’s multiple comparison test. (C) Statistical analysis performed on log-transformed data using Kruskal-Wallis test with Dunn’s multiple comparisons. (D, F) Statistical analysis represents Kruskal-Wallis test with Dunn’s multiple comparisons. LLOD = lower limit of detection; ns = not significant. * *p* < 0.05, ** *p* < 0.01, *** *p* < 0.001, **** *p* < 0.0001. Associated post-prime IgG and post-boost CD4^+^ and CD8^+^ T cell data in **Supplementary Figure S4**.

The H7 saRNA-NLC vaccine induced H7-specific systemic and mucosal immune responses in similar patterns to those induced by the H5 vaccine. Strong H7-binding serum IgG titers were consistently observed in animals receiving doses of at least 5 µg of H7 saRNA-NLC, regardless of administration route (**Figure 2B, Supplementary Figure S4A**). Immunization with 10 µg of H7 saRNA-NLC IN/IN was required to induce serum neutralization equivalent to immunization with 5 µg dosed IM/IN or IM/IM (**Figure 2C**), suggesting that any reduction in vaccine potency seen with IN administration, possibly due to lowering of effective dose delivered, may be overcome by a modest increase in nominal vaccine dose. The establishment of splenic antigen-reactive polyfunctional CD8^+^, polyfunctional CD4^+^, and IFNγ^+^ CD8^+^ T cells was likewise dose dependent, with comparable responses induced by IN/IN 10 µg, IM/IN 5 µg, and IM/IM 5 µg vaccination schedules (*p* > 0.05) (**Figure 2D, Supplementary Figure S4B-C**).

Induction of respiratory mucosal immunity following H7 saRNA-NLC vaccination was consistent with H5 saRNA-NLC vaccination results. Development of H7-binding IgA in BAL samples strictly necessitated at least one IN immunization (**Figure 2E**). Likewise, an IN/IN 10 µg dose established the highest mean frequency of lung-resident polyfunctional CD8^+^, polyfunctional CD4^+^, and IFNγ^+^ CD8^+^ T cells, while an IM/IN 5 µg dose established comparable responses at half the saRNA-NLC dose (*p* > 0.05) (**Figure 2F, Supplementary Figure S4D-E**). As with the H5 saRNA-NLC, no detectable mucosal IgA and low mucosal cellular immune responses were observed following IM-only administration of H7 saRNA-NLC. Together, these data demonstrate that our IN H7 saRNA-NLC vaccine formulation is immunogenic in mice, producing systemic and mucosal immune response patterns akin to those seen for the H5 saRNA-NLC vaccine, thereby validating the use of this saRNA-NLC vaccine platform for diverse influenza strains.

### Intramuscular administration of a bivalent H5/H7 saRNA-NLC vaccine induces strong systemic immunity without antigen competition but no detectable mucosal immunity

Having confirmed the immunogenicity of the monovalent H5 and H7 saRNA-NLC vaccines, we tested a bivalent H5/H7 saRNA-NLC vaccine formulation for vaccine compatibility and antigenic competition in the context of IM immunization. Mice were prime/boost vaccinated IM/IM with 5 µg of monovalent H5 or H7 saRNA-NLC vaccine, or 5 µg of each saRNA-NLC vaccine combined by simple mixing (10 µg saRNA total). Additional study groups were included to control for the total saRNA and NLC content of the vaccines by admixing 5 µg of each monovalent saRNA-NLC with 5 µg of the SEAP saRNA-NLC (**Supplementary Figure S5**). Systemic and mucosal humoral and cellular immune responses to both antigens were assessed as before after prime (**Supplementary Figures S6-S7**) and boost (**Figure 3, Supplementary Figures S6-S7**).

**Figure 3.**
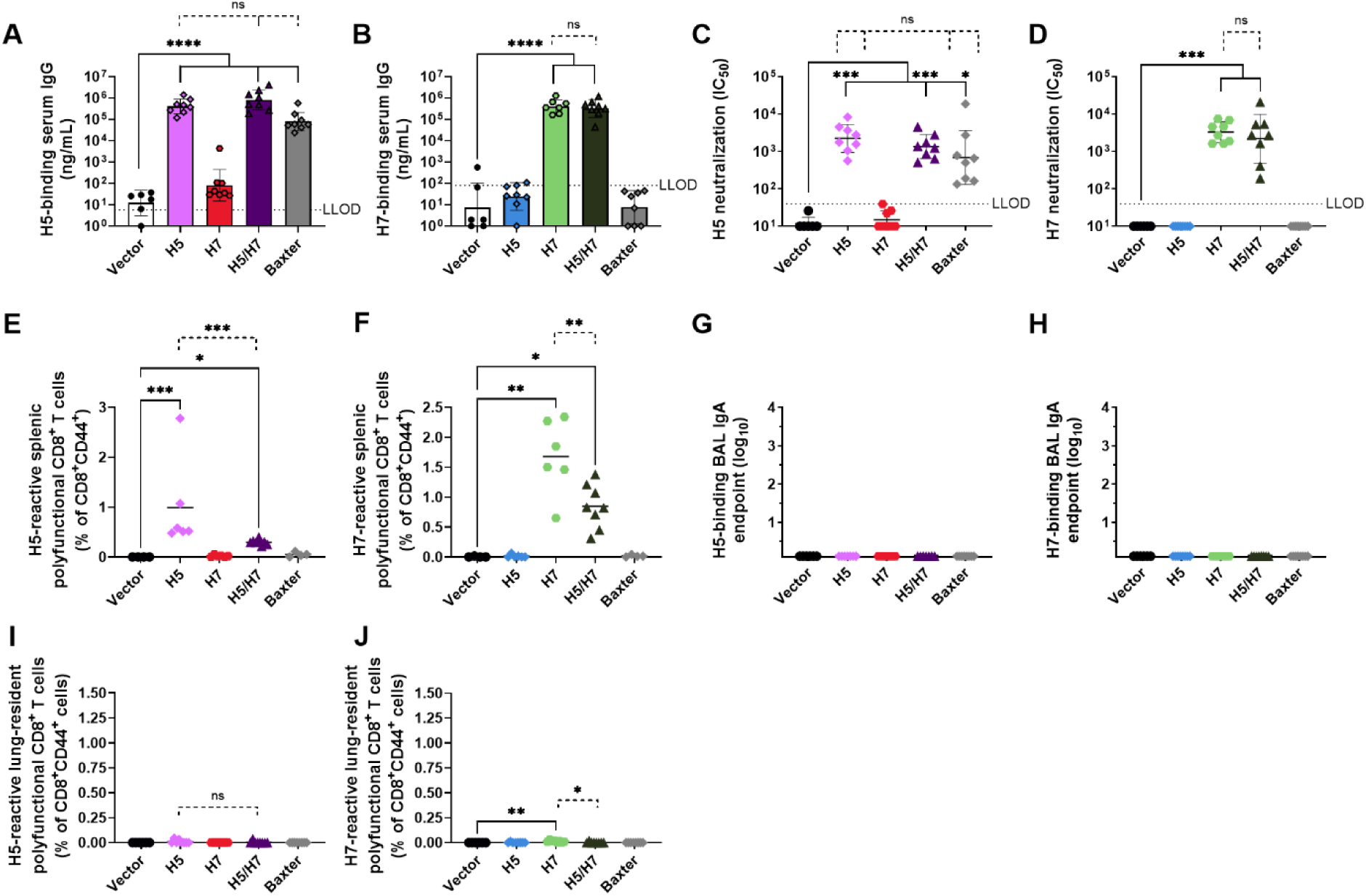
Intramuscular prime/boost dosing of bivalent H5/H7 saRNA-NLC establishes systemic immunity without antigen competition but no detectable mucosal immunity in mice. Post-boost serum (A) H5 and (B) H7-binding IgG titers, serum (C) H5 and (D) H7 pseudovirus neutralization capacity (IC_50_), splenic (E) H5- and (F) H7-reactive polyfunctional (IFN*γ*^+^ IL-2^+^ TNF*α*^+^) CD8^+^ T cells, (G) H5- and (H) H7-binding IgA titers in BAL samples, and lung-resident (CD69^+^ CD103^+^) (I) H5- and (J) H7-reactive polyfunctional (IFN*γ*^+^ IL-2^+^ TNF*α*^+^) CD8^+^ T cells. For all assays, *n* = 6 animals for vector control group, *n* = 8 for experimental groups. All groups were sex balanced. Two statistical hypotheses were tested within each figure, unless specifically listed, with filled lines showing comparisons to the vector control and dotted lines representing tests between key experimental groups. (A-B, G-H) Statistical analyses performed on log-transformed data using one-way ANOVA with Šídák’s multiple comparison test. (C-D) Statistical analysis performed on log-transformed data using Kruskal-Wallis test with Dunn’s multiple comparisons. (E-F) Statistical analysis performed using Kruskal-Wallis test with Dunn’s multiple comparisons (filled lines) or unpaired t tests (dotted lines). (I-J) Statistical analysis performed using Kruskal-Wallis test with Dunn’s multiple comparisons (filled lines) or Mann-Whitney tests (dotted lines). LLOD = lower limit of detection; ns = not significant. * *p* < 0.05, ** *p* < 0.01, *** *p* < 0.001, **** *p* < 0.0001. Associated post-prime and post-boost data in **Supplementary Figures S6-S7**.

We observed strong antigen-specific serum IgG and nAb responses against H5 or H7 antigens for both monovalent and bivalent saRNA-NLC vaccines, with no indication of significant IgG cross-binding or cross-neutralization between the antigens (**Figure 3A-D, Supplementary Figure S6A-F**). Splenic cellular immune responses followed a similar pattern, with induction by both monovalent and bivalent saRNA-NLC vaccines at notably high frequencies with respect to the overall T cell population (**Figure 3E-F, Supplementary Figure S6G-L**). Consistent with prior IM administrations, a limited mucosal immune response was induced by any IM-delivered vaccine, as seen by undetectable IgA titers in BAL samples and low frequencies (<0.2%) of lung-resident antigen-reactive CD8^+^ T cells (**Figure 3G-J, Supplementary Figure S7**), underscoring the value of IN administration of saRNA-NLC vaccines for generating localized respiratory mucosal immunity.

### Intranasal administration of a bivalent H5/H7 saRNA-NLC vaccine provides strong systemic and mucosal immunity against both viral targets

We next sought to confirm whether an intranasally-delivered bivalent H5/H7 saRNA-NLC vaccine would elicit both systemic and mucosal immune responses against both viral targets. As such, we implemented an IN version of the IM bivalent study design and compared immunogenicity between bivalent and monovalent formulations and controls.

After prime/boost vaccination, we observed comparable humoral responses between the monovalent and bivalent saRNA-NLC vaccines in terms of serum H5- or H7-binding IgG titers and pseudovirus neutralization, with no evidence of IgG cross-binding or cross-neutralization (**Figure 4A-D, Supplementary Figure S8A-F**). Splenic H5-specific and H7-specific polyfunctional CD8^+^ T cell responses were also comparable between monovalent and bivalent vaccines (**Figure 4E-F, Supplementary Figure S8G-L**), and consistent with previous studies (**Figures 1D and 2D**). Like systemic responses, mucosal responses were comparable after monovalent or bivalent saRNA-NLC vaccination. Strong, antigen-specific IgA responses were seen in BAL samples (**Figure 4G-H, Supplementary Figure S9A-B**), and similar frequencies of lung-resident antigen-responsive bifunctional (IFNγ^+^ TNFα^+^) T cells were detected, all with little evidence of cross-reactivity or inhibition by bivalent dosing (**Figure 4I-J, Supplementary Figure S9C-H**). These data verify that an IN/IN prime/boost regimen of the bivalent H5/H7 saRNA-NLC vaccine stimulates excellent systemic and mucosal immunity without antigen interference or competition.

**Figure 4.**
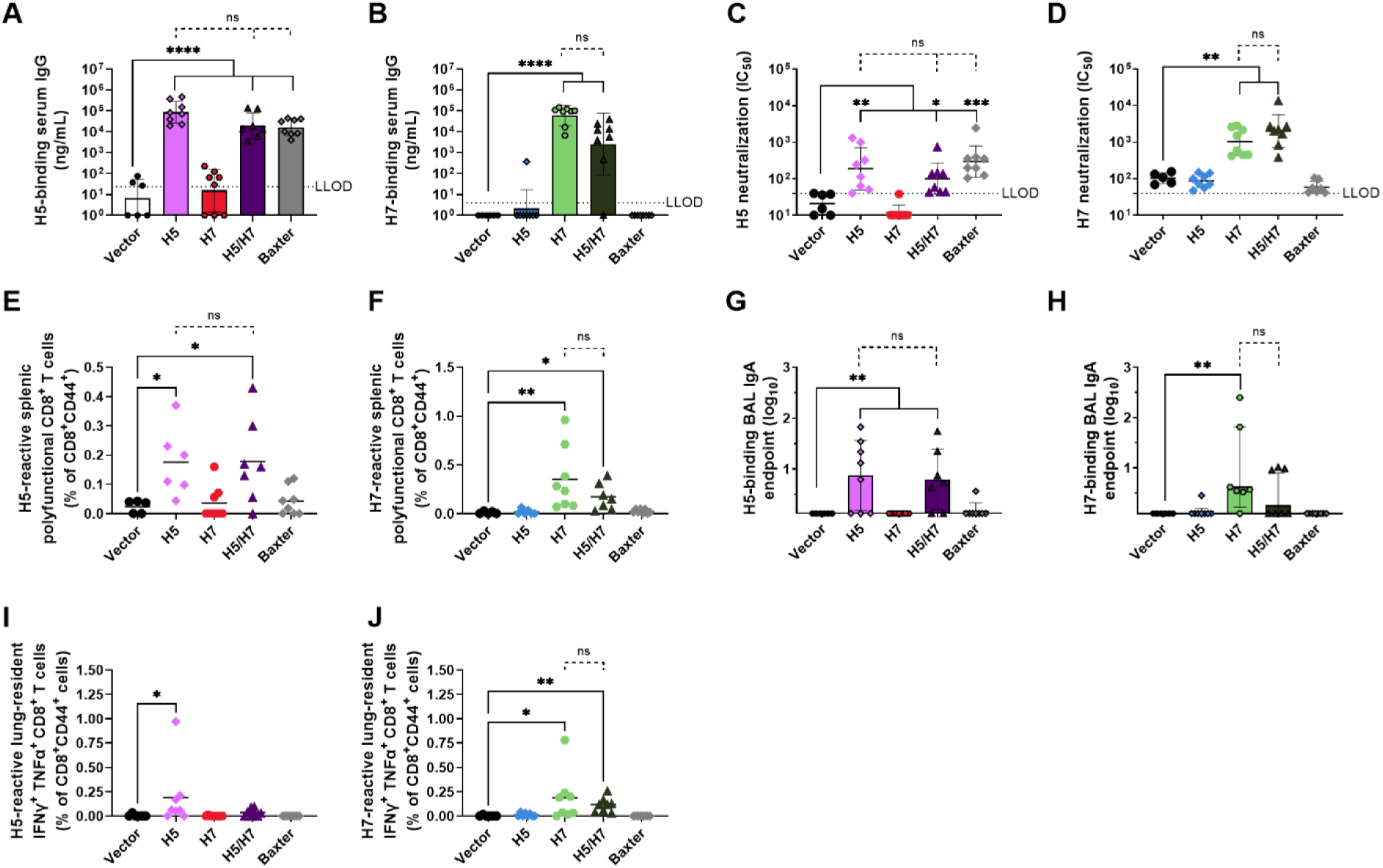
Intranasal vaccination with bivalent H5/H7 saRNA-NLC establishes both systemic and mucosal immunity in mice. Post-boost serum (A) H5- and (B) H7-binding IgG titers, serum (C) H5 and (D) H7 pseudovirus neutralization capacity (IC_50_), splenic (E) H5- and (F) H7-reactive polyfunctional (IFN *γ*^+^ IL-2^+^ TNF*α*^+^) CD8^+^ T cells, (G) H5- and (H) H7-binding IgA antibody titers in BAL samples, and lung-resident (CD69^+^ CD103^+^) (I) H5- and (J) H7-reactive bifunctional (IFN*γ*^+^ TNF*α*^+^) CD8^+^ T cells. For all assays, *n* = 6 animals for vector control group, *n* = 8 for experimental groups. All groups were sex balanced. Two statistical hypotheses were tested within each figure, unless specifically listed, with filled lines showing comparisons to the vector control and dotted lines representing tests between key experimental groups. (A-B, G-H) Statistical analyses performed on log-transformed data used one-way ANOVA with Šídák’s multiple comparison test. (C-D) Statistical analysis performed on log-transformed data used Kruskal-Wallis test with Dunn’s multiple comparisons. (E-F) Statistical analysis performed using Kruskal-Wallis test with Dunn’s multiple comparisons (filled lines) or unpaired *t* tests (dotted lines). (I-J) Statistical analysis performed using Kruskal-Wallis test with Dunn’s multiple comparisons (filled lines) or Mann-Whitney tests (dotted lines). LLOD = lower limit of detection; ns = not significant. * *p* < 0.05, ** *p* < 0.01, *** *p* < 0.001, **** *p* < 0.0001. Associated post-prime and post-boost data in **Supplementary Figures S8-S9**.

### Route of vaccination of a bivalent H5/H7 HA saRNA-NLC vaccine influences mucosal but not systemic immunogenicity

Bivalent intramuscularly and intranasally administered saRNA-NLC vaccines were observed to establish consistent systemic immune responses in mice; however, there were notable differences in the establishment of mucosal immunity between IM and IN vaccination. To optimize the induction of systemic and mucosal immunity by our vaccine, we tested permutations of heterologous vaccination regimens. Mice were vaccinated with 10 µg of the bivalent H5/H7 saRNA vaccine (5 µg of each saRNA) either through homologous (IM/IM and IN/IN) or heterologous (IM/IN and IN/IM) routes, and immune responses were assayed as above.

Consistent with the other monovalent and bivalent dosing studies, the route of administration did not result in significant differences in the systemic antibody response against either antigen after prime/boost vaccination (**Figure 5A-D, Supplementary Figure S10A-F**). Mucosal immunity, both humoral and cellular, still required IN vaccination either as part of a homologous or a heterologous vaccination regimen (**Figure 5E-F, Supplementary Figure S11A-B**). The IM/IN combination appeared to induce an optimal combination of both systemic and mucosal immune responses. Interestingly, the IN/IM combination resulted in higher spread or numbers of non-responders in systemic nAbs, IgA in BAL samples, and cellular responses (**Figure 5, Supplementary Figures S10-S11**). Thus, we proceeded to test IM/IN immunization further in efficacy models alongside the homologous IN/IN dosed vaccine candidate.

**Figure 5.**
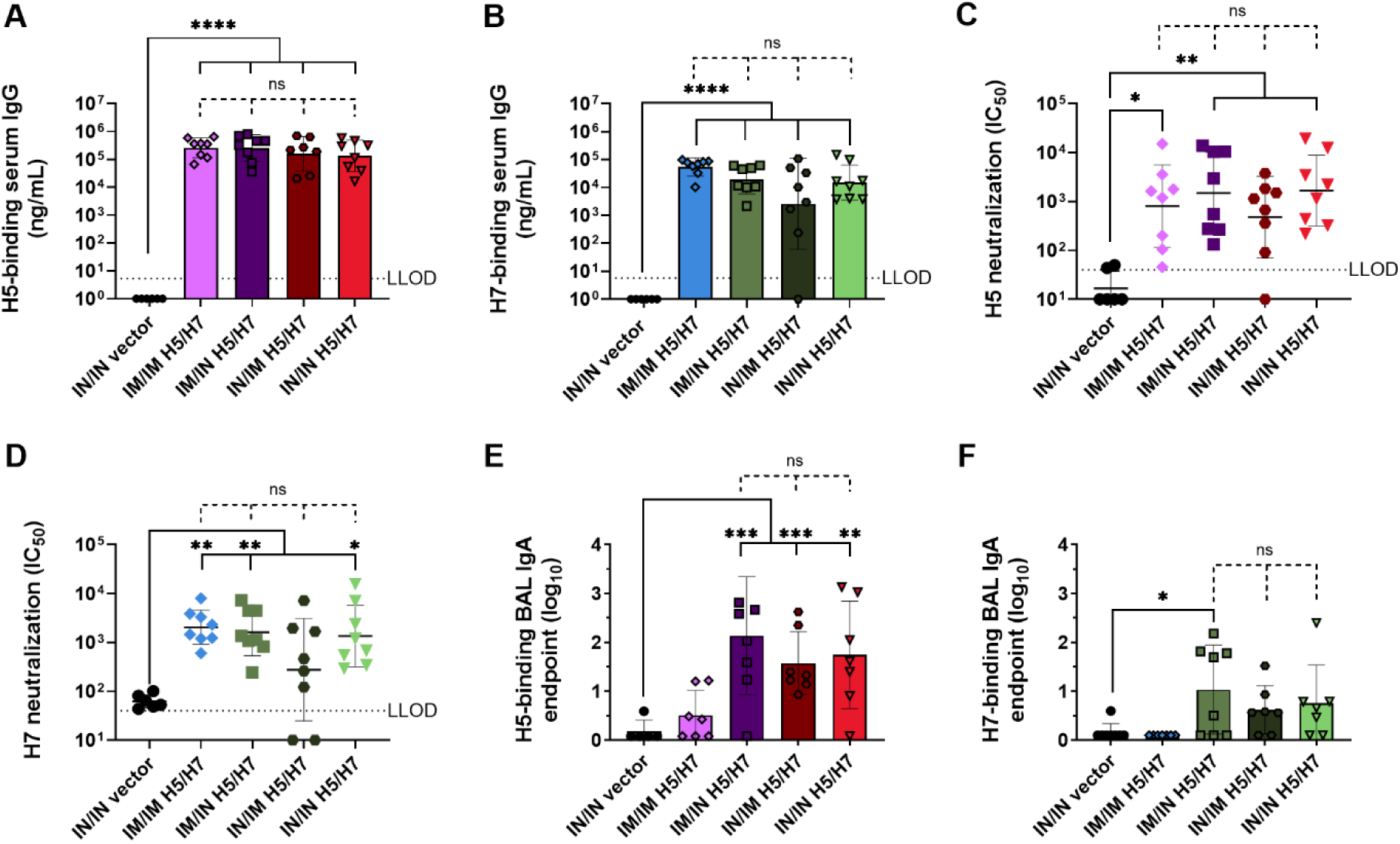
Homologous and heterologous routes of bivalent vaccination establish comparable systemic immunogenicity but differential mucosal immunogenicity in mice. Post-boost serum (A) H5 and (B) H7 HA-specific IgG titers, serum (C) H5 and (D) H7 pseudovirus neutralization capacity (IC_50_), and (E) H5- and (F) H7-binding IgA antibody titers in BAL samples. For all assays, *n* = 6 animals for vector control group, *n* = 8 for experimental groups. All groups were sex balanced. (A-B) Statistical analyses performed on log-transformed data using one-way ANOVA with Šídák’s multiple comparison test. (C-D) Statistical analysis performed on log-transformed data used Kruskal-Wallis test with Dunn’s multiple comparisons. (E-F) Statistical analysis represents one-way ANOVA with Šídák’s multiple comparison test. Two statistical hypotheses were tested within each figure, with filled lines showing comparisons to the vector control and dotted lines representing tests between key experimental groups. LLOD = lower limit of detection; ns = not significant. * *p* < 0.05, ** *p* < 0.01, *** *p* < 0.001, **** *p* < 0.0001. Associated post-prime and post-boost data in **Supplementary Figures S10-S11**.

### Monovalent H7 and bivalent H5/H7 saRNA-NLC vaccines protect ferrets from virus-induced morbidity and promote rapid viral clearance in a nonlethal H7N9 ferret challenge model

We tested the protective efficacy of our IN/IN and IM/IN delivered H7 saRNA-NLC vaccine candidates in an established sublethal high-dose H7N9 ferret challenge model. Sex-balanced groups of Fitch ferrets, *n* = 6 per group, were vaccinated either IN/IN or IM/IN with either 25 µg of monovalent H7 saRNA-NLC or 50 µg (25 µg of each antigen) of bivalent H5/H7 saRNA-NLC. Control groups included a prime/boost IN/IN dose of 50 µg SEAP saRNA-NLC vector control or a prime/boost dose of 7.5 µg of H7 HA from the National Institute for Biological Standards and Control (NIBSC) unadjuvanted influenza antigen A/Anhui/1/2013 (NIBRG-268, NIBSC code 14/250) delivered IM in 250 µL. Three weeks after boost vaccination, ferrets were challenged intranasally with 1 x 10^6^ pfu of A/Anhui/2013 virus and observed for 14 days with regular body weight measurements, scoring of clinical disease, and nasal swabs for viral load measurements.

All animals survived throughout the 14-day challenge period (**Figure 6A**), as expected for this non-lethal challenge model. Only vector control immunized animals developed higher average body temperatures from Days 3 to 5 post-challenge (**Supplementary Figure S12**) and brief clinical signs of disease in a subset of animals noted 8 days post-challenge, which quickly resolved and was not seen in any H7 vaccinated animal (**Figure 6B**). All H7 vaccinated animals were protected against weight loss relative to vector control immunized animals (**Figure 6C**). Analysis of viral titers in nasal wash samples collected after challenge demonstrated that, while most animals across all study groups had similar viral loads 1 and 3 days post-challenge, reflecting equivalent challenge doses, by Day 5, viral loads in all H7 vaccinated animals were reduced to undetectable levels (**Figure 6D-F**).

**Figure 6.**
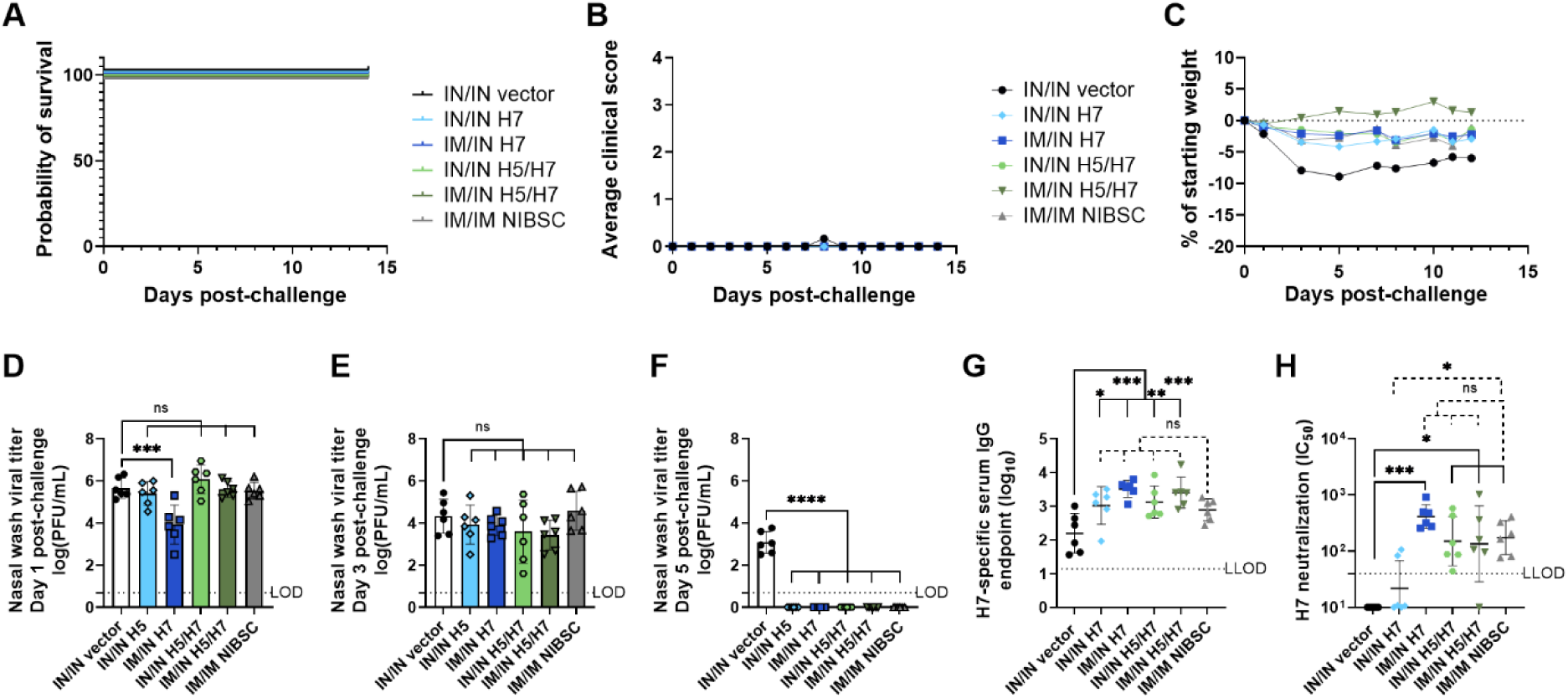
Intranasal and heterologous routes of monovalent H7 and bivalent H5/H7 saRNA-NLC vaccination protect ferrets from sublethal H7N9 challenge. For all assays, *n* = 6 (3 male and 3 female) animals per group. (A) Animal survival after challenge with 1 x 10^6^ pfu of intranasally-delivered A/Anhui/2013. (B) Averaged clinical scores. (C) Percent change in body weight from date of challenge. Two statistical hypotheses were tested within each figure, with filled lines showing comparisons to the vector control and dotted lines representing tests between key experimental groups. (D-F) Viral load in nasal wash at Day (D) 1, (E) 3, and (F) 5 post-challenge. Statistical analysis performed on log-transformed data used one-way ANOVA with Holm-Šídák’s multiple comparison test. (G) H7-binding IgG antibody titers in pre-challenge ferret serum. (H) H7 pseudovirus neutralization capacity (IC_50_) of pre-challenge ferret serum. (G) Statistical analysis performed on log-transformed data used one-way ANOVA with Dunnett’s multiple comparisons test. (H) Statistical analysis performed on log-transformed data using the Kruskal-Wallis test with Dunn’s multiple comparisons. LLOD = lower limit of detection; NIBSC = NIBSC Influenza Antigen A/Anhui/1/2013 (H7N9); ns = not significant. * *p* < 0.05, ** *p* < 0.01, *** *p* < 0.001, **** *p* < 0.0001. Associated body temperature data in **Supplementary Figure S12**.

Post-boost, pre-challenge serology indicated that all saRNA-NLC vaccine regimens induced H7-binding IgG responses measurably above the vector control group, with signal in the control group attributed to non-specific background signal (**Figure 6G**). Of note, serum IgG and nAb titers induced by the saRNA-NLC vaccines comparable to those induced by the NIBSC H7N9 antigen (**Figure 6G-H**). This challenge study thus demonstrated that 25 µg prime/boost doses of H7 or H5/H7 saRNA-NLC vaccines, regardless of IN/IN or IM/IN administration, successfully protect ferrets from H7N9 challenge-induced morbidity.

### Monovalent H5 and bivalent H5/H7 saRNA-NLC vaccines completely protect ferrets from lethal H5N1 challenge

We then tested the protective efficacy of our IN/IN and IM/IN delivered H5 saRNA-NLC vaccine candidates in an established lethal high-dose H5N1 ferret challenge model. Sex-balanced groups of Fitch ferrets, *n* = 6 per group, were vaccinated either IN/IN or IM/IN with either 25 µg of monovalent H5 saRNA-NLC or 50 µg (25 µg of each antigen) of bivalent H5/H7 saRNA-NLC. Control groups included a prime/boost IN/IN dose of 50 µg SEAP saRNA-NLC vector control, or a prime/boost dose of 7.5 µg alum-adjuvanted Baxter H5N1 vaccine delivered IM. Three weeks after boost vaccination, ferrets were challenged intranasally with 1 x 10^6^ pfu of A/Vietnam/1203/2004 virus and observed for 14 days with regular body weight measurements, scoring of clinical disease, and nasal swabs for viral load measurements.

All animals in the vector control saRNA-NLC treated group succumbed to infection by 1 week post-challenge (**Figure 7A**). One animal in the Baxter vaccinated group succumbed to infection 9 days after challenge. All animals vaccinated with H5 or H5/H7 saRNA-NLC vaccines were fully protected from virus-induced mortality and from the development of clinical symptoms, weight loss, and severe fever (**Figure 7A-C, Supplementary Figure S13A**). Analysis of viral titers in nasal wash samples collected after challenge demonstrated that, while all animals had similar viral loads 1 day after infection, reflecting equivalent challenge doses, viral loads in all H5 vaccinated animals were significantly reduced by 3 days post-challenge relative to the vector control treated animals (**Figure 7D-E**). Strikingly, no virus was detectable in the nasal wash of H5 vaccinated animals 5 days post-challenge, whereas vector control animals displayed viral titers equivalent to those 1 day post-challenge (**Figure 7F**). Lung viral titers were evaluated upon necropsy after the animals either succumbed to infection or were euthanized, and these indicated that all deaths in the study were virus associated (**Supplementary Figure S13B-C**).

**Figure 7.**
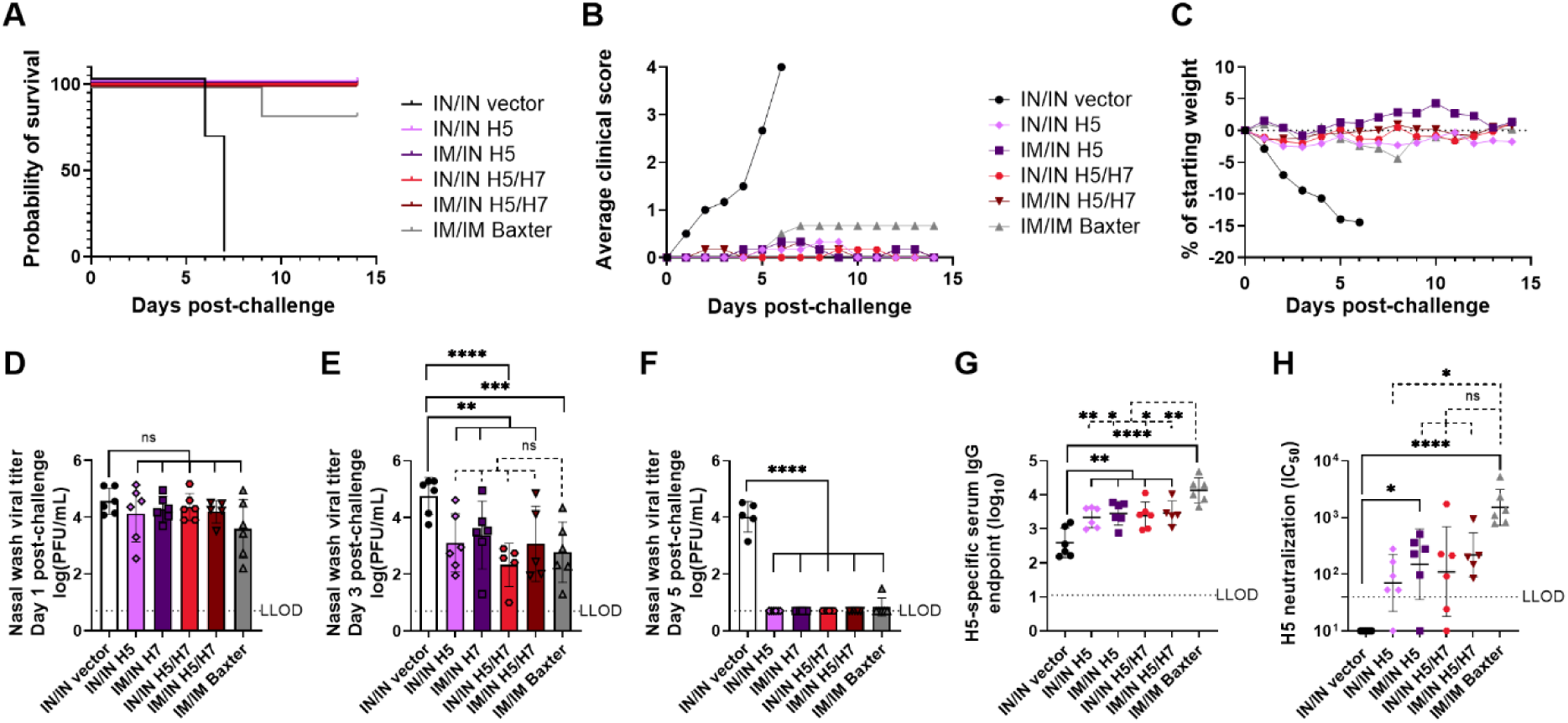
Intranasal and heterologous routes of monovalent H5 and bivalent H5/H7 saRNA-NLC vaccination protect ferrets from lethal H5N1 challenge. For all assays, *n* = 6 (3 male and 3 female) animals per group. (A) Animal survival after challenge with 1 x 10^6^ pfu of intranasally-delivered A/Vietnam/1203/2004. (B) Averaged clinical scores. (C) Percent change in body weight from date of challenge. Two statistical hypotheses were tested within each figure, with filled lines showing comparisons to the vector control and dotted lines representing tests between key experimental groups. (D-F) Viral load in nasal wash at Day (D) 1, (E) 3, and (F) 5 post-challenge. Statistical analysis performed on log-transformed data used one-way ANOVA with Holm-Šídák’s multiple comparison test. (G) H5-binding IgG antibody titers in pre-challenge ferret serum. (H) H5 pseudovirus neutralization capacity (IC_50_) of pre-challenge ferret serum. (G) Statistical analysis performed on log-transformed data used one-way ANOVA with Dunnett’s multiple comparisons test. (H) Statistical analysis performed on log-transformed data using the Kruskal-Wallis test with Dunn’s multiple comparisons. LLOD = lower limit of detection; ns = not significant. * *p* < 0.05, ** *p* < 0.01, *** *p* < 0.001, **** *p* < 0.0001.

Post-boost, pre-challenge serology indicated the four saRNA-NLC vaccine regimens successfully induced H5-binding IgG titers measurably above the vector control group, with signal in the control group attributed to non-specific background signal (**Figure 7G**). Of note, the Baxter vaccine induced trending or significantly higher levels of H5-binding IgG and nAbs compared to the four IN/IN or IM/IN saRNA-NLC vaccine groups (**Figure 7G-H**). Despite saRNA-NLC vaccines inducing a lower humoral response than the Baxter vaccine, saRNA-NLC vaccines were completely protective, suggesting that serum antibody titers are insufficient correlates of protection for mucosal H5N1 influenza vaccines. In sum, this challenge study demonstrated that 25 µg prime/boost doses of the H5 or H5/H7 saRNA-NLC vaccines, regardless of IN/IN or IM/IN administration, fully protect ferrets from H5N1 viral challenge-induced morbidity and mortality.

## DISCUSSION/CONCLUSION

The studies presented here demonstrate the immunogenicity and efficacy of IN- and IM/IN-administered saRNA-NLC vaccines against two different pre-pandemic influenza strains. Vaccination with monovalent and bivalent saRNA-NLC formulations induced robust systemic immune responses in mice against the cognate virus, with no signs of immune competition between antigens after bivalent dosing. Intranasal administration, either as a prime/boost or as a boost of a previous IM vaccination, added key mucosal IgA and lung-resident cellular immune responses that were not induced by IM injection of the saRNA-NLC vaccines or the previously-authorized adjuvanted inactivated whole virion H5N1 Baxter vaccine. In ferrets, the H5 saRNA-NLC vaccines demonstrated 100% efficacy, protecting animals against viral morbidity and mortality regardless of vaccine valency or route of administration.

Our results underscore the importance of administration route on the nature of immune response. Like other influenza vaccines, these saRNA-NLC vaccines induced systemic humoral and cellular immune responses (*35–37*). However, the mucosal immune responses to the intranasally administered saRNA-NLC vaccines are unique relative to approved intramuscularly administered vaccines. When delivered intranasally, these saRNA-NLC vaccines established robust local respiratory mucosal immune responses not seen following IM administration of either saRNA-NLC or other influenza vaccines. The mucosal responses observed here, namely IgA in BAL samples and lung-resident (CD69^+^ and/or CD103^+^) antigen-specific CD4^+^ and CD8^+^ T cells, are associated with enhanced vaccine efficacy, control of early infection and transmission, and improved outcomes in the context of influenza infection (*37–40*).

Several mRNA influenza vaccine candidates have been developed previously (*41–43*); however, none has yet been demonstrated to support IN vaccination. This deficit may explain the inability of licensed SARS-CoV-2 mRNA vaccines to prevent mild infection in respiratory tissues (*20, 21*), possibly due to lack of established mucosal immunity (*22*). The flexible mode of delivery demonstrated by this saRNA-NLC vaccine platform is likely due to the unique properties of the NLC, which provides excellent RNA delivery while maintaining low reactogenicity (*31*). This ability to tune the immune profile by changing route of vaccine administration is a major advantage of this saRNA-NLC platform as a pandemic response tool.

Notably, despite inducing lower serum IgG and nAb titers in ferrets relative to the intramuscularly delivered adjuvanted, inactivated whole virion vaccine, the H5-expressing saRNA-NLC vaccines provide stellar protection of animals from H5N1 challenge. This equivalent or improved protection of the saRNA-NLC vaccines despite lower serum antibody titers suggests that for mucosal influenza vaccines, serum antibody titers are insufficient correlates of protection. This is in contrast with the well-established standard of a 1:40 HA inhibition assay readout as an indicator of seroconversion and protection for traditional intramuscularly administered influenza vaccines. Mucosal correlates of protection have been difficult to establish to date due to the difficulty of sample collection and determination of protective mucosal assay thresholds (*44, 45*). These challenges pose a barrier to optimal design of human clinical trials for mucosal vaccines. For this reason, further work examining potential mucosal correlates of protection in primate models is planned.

Despite the success of licensed IIVs and LAIVs, aspects of their manufacturing processes, such as reliance on egg- or cell-based virus production, are slow and can impair yearly efficacy, imposing limitations on successful seasonal and pandemic influenza response (*46*). Antigenic drift over the 6-month lead time required for production and distribution can result in seasonal vaccine strain mismatch and an inability to rapidly develop vaccines for emerging strains of pandemic potential (*47*). In contrast, RNA vaccine platforms combine agile modification of antigen sequences with ease and rapidity of *in vitro* transcription-based manufacturing to allow for manufacture of vaccines targeting emerging strains in a fraction of the time required for IIVs and LAIVs (*48*), as needed for pandemic preparedness and response. The saRNA-NLC system is particularly amenable to rapid manufacturing and nimble adaptation to emerging viral strains. Unlike that of lipid nanoparticles used for other RNA vaccines, manufacture of the NLC RNA delivery component is straightforward and easily scalable, using standard emulsification equipment in place at most vaccine manufacturers worldwide. The NLC is additionally stable for long periods (>1 year at refrigerated conditions) in the absence of RNA (*31*), allowing for component stockpiling and last-minute changes in the identity of the RNA with which it is complexed in order to reflect circulating viral strains in real time.

While carefully designed and executed, these studies do have limitations. Pipet delivery of IN vaccines likely results in variability in delivered dose, which may contribute to the heterogeneity in immune response seen particularly for intranasally dosed animals. Our H5N1-targeting saRNA-NLC vaccine expresses HA from the prototypic A/Vietnam/1203/2004 H5N1 influenza strain, rather than any of the panzootic highly-pathogenic avian H5N1 clade 2.3.4.4b strains circulating in US dairy cattle in 2024 (*3*). This antigen selection was deliberate, intended to establish this IN RNA vaccine technology with existing reagents and animal efficacy models, with the knowledge that RNA vaccines allow for rapid reformulation against emerging antigen targets when required. While assessment of these saRNA-NLC vaccines’ cross-protection against currently-circulating strains was not a part of this study, recent studies of cross-neutralizing antibodies induced by licensed H5N1 vaccines suggest some cross-protection against the current circulating strains may be expected, which may possibly be enhanced by respiratory mucosal immunity provided by IN dosing (*49, 50*). Finally, an additional weakness of the work presented here is that many of the conclusions from this study rely on the immune readouts after IN vaccine administration in mice and ferrets. As both of these animal models have substantially different respiratory structures and volumes than humans, this may result in differential vaccine material distribution relative to what would be seen in humans, emphasizing the need for testing in a non-human primate (NHP) model.

Future directions include toxicity testing and rapid progression into a Phase 1 clinical trial to evaluate safety and immunogenicity of these intranasally delivered saRNA-NLC vaccines in humans. Simultaneously, detailed immunogenicity and efficacy studies in NHPs are planned, with immunological profiling to elucidate the mode of action of these intranasally delivered vaccines in animals with similar nasal anatomy and immune responses to those of humans. A desired secondary outcome of this study is the identification of potential mucosal correlates of protection, as suggested by our ferret study outcomes. Similarly, compatibility and characterization studies of this product in nasal spray devices is necessary for enabling mid- and late-stage clinical trials and future vaccine commercialization.

This vaccine platform is a powerful potential tool for pandemic preparedness and response efforts. Amenable to multiple routes of parenteral administration with an identical formulation, the saRNA-NLC system can be flexibly used to combat pathogens of different natures (respiratory vs. mosquito borne, etc.). Furthermore, when lyophilized, saRNA-NLC vaccines are stable for at least 2 years at non-frozen temperatures (*31, 32*), thus demonstrating improved thermostability over other RNA vaccines and negating requirements for deep cold-chain storage and complex distribution (*19, 51*). Finally, the recent US FDA greenlighting of self-administration of the IN FluMist vaccine (*52*) opens the door to a potential thermostable, self-administrable saRNA vaccine as a pandemic response tool. If demonstrated to be safe and immunogenic in humans, these IN saRNA-NLC influenza vaccines would represent a first-in-class technology with key advantages over current influenza vaccines in terms of flexible mode of delivery, mucosal immune stimulation, agile antigen targeting, and desirable product profile. Such a product, with long-term thermostability and self-administration potential, could represent an invaluable tool in pandemic preparedness and response.

## MATERIALS AND METHODS

### Experimental design

Details on sample size, timepoints, data inclusion, and selection of endpoints are listed in the animal-specific study protocols below. Group sizes, listed for each study, were based off power calculations that allowed for differentiation of dosing and formulation effects in immunogenicity assays below. Assays were run with technical replicates and internal controls to control for intra- and inter-assay variability. No exclusion or outlier criteria were utilized for these study data. Studies were not blinded.

### Development-grade saRNA production

HA-expressing saRNA template plasmids were created by subcloning gene blocks containing the influenza H5N1 HA (GenBank ID EU122404.1) or H7N9 HA (NCBI Reference Sequence YP_009118475.1) antigens into a plasmid containing a 5’ UTR, a 3’ UTR, and non-structural proteins derived from the attenuated TC83 strain of VEEV. saRNA was then produced using an *in vitro* transcription protocol and linearized DNA templates as previously described (*31, 32, 53*). RNA was purified via AKTA using CaptoCore700 resin (Cytiva) then buffer exchanged and concentrated using a 500 kDa cut-off mPES hollow fiber tangential flow filter membrane (Repligen). For small-scale lots, RNA was lithium chloride precipitated followed by ethanol precipitation to remove proteins and residual reagents. Concentration and yield were determined using spectrophotometry and validated by RiboGreen assay. RNA integrity was validated by gel electrophoresis.

### Development-grade NLC production

The NLC was formulated as previously described (*31, 32, 53*) as a solid/liquid oil core stabilized in an aqueous buffer by surfactants. Squalene (Sigma), sorbitan monostearate (Span 60; Spectrum Chemical Mfg. Corp.), DOTAP (N-[1-(2,3-dioleoyloxy)propyl]-N,N,N-trimethylammonium chloride; Lipoid), and glyceryl trimyristate (Dynasan 114; Sigma) were emulsified with a 10 mM sodium citrate trihydrate (Spectrum Chemical Mfg. Corp.) buffer-polysorbate 80 (Tween 80; Millipore) aqueous phase at 7,000 rpm using a high-speed laboratory emulsifier (Silverson Machines) followed by high-shear homogenization at 30,000 psi in a microfluidizer (Microfluidics M-110P). The end product was terminally filtered with a 0.22 µm polyethersulfone filter and stored at 2-8°C.

### Vaccine complexing and characterization

Monovalent vaccine and SEAP vector control material were prepared by diluting the saRNA to 2X the final concentration of the complex, in a background of 10 mM sodium citrate pH 6.3 and 10% w/v sucrose. The NLC stock was similarly diluted to 2X the final NLC concentration in the same background buffer. These diluted components were combined 1:1 by volume via pipette mixing. The complexes were then incubated on ice for 30 minutes before storage at -80°C. Complexed material was prepared at a nitrogen:phosphate (N:P) ratio of 8-10, indicating the ratio of nitrogen in the DOTAP component of the NLC to phosphate in the RNA backbone. Bivalent vaccines were prepared by simple pipette mixing of each monovalent vaccine 1:1 by volume.

The hydrodynamic diameter of the saRNA-NLC complexes was determined by dynamic light scattering of 1:100 diluted sample in water using a Zetasizer Pro (Malvern) with a backscatter angle of 173°. Measurements were carried out at 25°C with a dispersant and sample viscosity value of 0.8872 cP (water). Three repeat measurements per sample were collected and averaged.

Agarose gel electrophoresis was used to evaluate the saRNA size and integrity after complexing. Complexed vaccine samples and saRNA only controls were diluted to 40 ng/µL saRNA in nuclease-free water, then extracted using phenol:chloroform:isoamyl alcohol (25:24:1). Each extracted sample was mixed with NorthernMax-Gly Sample Loading Dye (Invitrogen), heated, and loaded onto a 1 cm thick, 1% (w/v) agarose gel. The gels were run at 120 V for 45 minutes and imaged under UV lighting (BioRad ChemiDoc MP Imaging System). Gels contained an RNA ladder (Ambion) for size reference.

### Mouse studies

Mouse studies were performed in accordance with national (NIH *Guide for the Care and Use of Laboratory Animals*) and institutional guidelines and approved by the Bloodworks Northwest Research Institute’s Institutional Animal Care and Use Committee (IACUC protocol 5389-01).

C57BL/6J mice (*n* = 8 per vaccine group, 6 per vector control group) sex balanced and between 6 and 8 weeks of age at study onset were obtained from The Jackson Laboratory (Harbor, ME). Body weights were collected up to 4 days post-vaccination to assess adverse effects. Animals were prime/boost immunized either IN or IM 3 weeks between doses, and blood was collected retro-orbitally on isoflurane-sedated mice. Animals were euthanized using CO_2_ followed by cervical dislocation in accordance with the recommendation of the American Veterinary Medical Association, followed by collection of BAL, lung, and spleen tissues.

Data were collected at pre-defined weekly timepoints. Euthanasia criteria (20% weight loss, moribund state) were not met by any animals during the studies. Prospective data inclusion/exclusion criteria were based on animal condition before, during, and after vaccination; vector and/or naïve animal controls; and internal assay standards. Each study was conducted once, with key experimental group conditions repeated in subsequent studies.

### Preparation of murine tissues into single-cell suspensions

#### Spleens

Spleens were processed as previously described (*32*). Briefly, spleens were individually homogenized by manual maceration through a 70 µm cell strainer. Red blood cells were lysed for 1 minute with ACK Lysing Buffer (ThermoFisher) and quenched with RPMI medium. Samples were filtered through a 2 mL Acroprep filter plate and resuspended in RPMI medium with 10% FBS, and 1-2 x 10^6^ cells/well were seeded in 96-well round-bottom plates in preparation for peptide stimulation.

#### Lungs

Lungs were processed as previously described (*32*). Briefly, lung tissue was dissociated via enzymatic digestion using Hanks’ Balanced Salt Solution supplemented with 10% Liberase (MilliporeSigma), 10% aminoguanidine, 0.1% KN-62, and 1.25% DNase. Lungs and enzymatic mix were added to a gentleMACS M tube (Miltenyi Biotec), run on a lung dissociation program in a gentleMACS Dissociator, incubated at 37°C and 5% CO_2_ for 30 minutes, then run again on the gentleMACS Dissociator. The resulting slurry was washed with RPMI and then filtered through a 70 µm MACS SmartStrainer (Miltenyi Biotec). Cells were counted using a Guava easyCyte cytometer before plating in 96-well round-bottom plates at 1 x 10^6^ cells/well in RPMI medium with 10% FBS in preparation for peptide stimulation.

### Stimulation and intracellular staining of murine single-cell suspensions

H5 peptides (JPT Peptide Technologies #PM-INFA-HAIndo) used for stimulation comprised a pool of 140 peptides (15mers, 11 amino acid overlaps) derived from the full-length HA of influenza A/Indonesia/DCD835/2006 (H5N1), which has a 96.64% sequence homology to the H5 HA from A/Vietnam/1203/2004 used in these studies. H7 peptides (custom ordered from GenScript) comprised a pool of 138 peptides (15mers, 11 amino acid overlaps) derived from the full-length HA sequence from A/Anhui/2013.

Stimulation mixes consisted of RPMI media, 10% FBS, 50 μM 2-mercaptoethanol, α-CD28 (BD), brefeldin A (BioLegend), and of three stimulation treatments: 0.26% dimethyl sulfoxide (DMSO) as a negative stimulation control, 0.2 µg/well (1 µg/mL) per peptide of H5 or H7 peptide pools in 0.26% DMSO, or phorbol myristate acetate-ionomycin positive stimulation solution.

Spleen and lung lymphocytes were stained for viability with Zombie Green (BioLegend #423112), then Fc receptors were blocked with CD16/CD32 antibody (Invitrogen #14-0161-82). Cells were surface stained for mouse CD4 (APC-Cy7, BD Bioscience #552051), CD8 (BV510, BD Biosciences #563068), CD44 (PE-CF594, BD Biosciences #562464), and CD154 (BV605, BD Bioscience #745242). Lung cells were additionally stained for mouse CD69 (PE, BD Biosciences #561932) and CD103 (BV711, Biolegend #121435). After extracellular staining, cells were fixed and permeabilized using BD Cytofix/Cytoperm (BD Biosciences #554714) and stained for intracellular cytokines with mouse TNFα (BV421, BioLegend #506328), IL-2 (PE-Cy5, BioLegend #503824), IFNγ (PE-Cy7, BD Biosciences #557649), IL-5 (APC, Biolegend # 504306), and IL-17A (AF700, BD Biosciences #560820). Cells were run on a CytoFLEX flow cytometer (Beckton Dickson) and analyzed with FlowJo v.10 (BD Biosciences). Representative gating strategies are shown in **Supplementary Figures S14-S15**. Cells triple positive for TNFα, IL-2, and IFNγ were considered polyfunctional T cells. Of note, flow cytometry data in Figure 4 excluded IL-2 (i.e., bifunctional instead of polyfunctional) due to poor data quality based on in-run standards. Samples with insufficient threshold events (<50,000) collected per well and low viability (<40% Zombie Green negative) were excluded from analysis.

### Ferret efficacy study

Ferret studies were performed in accordance with national (NIH *Guide for the Care and Use of Laboratory Animals*) and institutional guidelines for animal care of laboratory animals and approved by the Colorado State University IACUC (protocol 4228) and were conducted as previously described (*54*). Fitch ferrets (*Mustela putorius fero*) were purchased from Triple F Farms (Gillett, PA) at approximately 3 months of age and confirmed to be serologically negative for influenza virus by microneutralization testing with H5N1-PR8 or H7N9 recombinant influenza virus prior to initiating immunization; sera for this testing were obtained prior to challenge by jugular venipuncture under ketamine-xylazine anesthesia. Vaccination with saRNA-NLC complexes consisted of a prime dose of 25 µg given either IM or IN to a cohort of 6 ferrets per group. A subsequent boost dose of 25 µg saRNA was delivered IN 21 days post-prime. The alum-adjuvanted inactivated whole virion H5N1 vaccine (“Baxter;” BEI Resources, NIAID, NIH: A/H5N1 Influenza Vaccine, Inactivated Whole Virion (A/Vietnam/1203/2004), Vero-Cell Derived, Adjuvanted, 15 Micrograms HA, NR-12143) was used as a positive control, while SEAP saRNA was used as a negative control. Ferrets were regularly observed for symptoms and clinical scores after each vaccination and during the challenge phase. Clinical scores were defined as 0 = normal; 1 = possibly less active than normal OR not grooming as normal (questionably sick or mild illness); 2 = less active than normal when viewed in cage but appears nearly normal and responsive when handled (sick); 3 = reluctance to rise when viewed in cage and not normally responsive when handled by humans OR weight loss up to 15% of pre-challenge value; 4 = failure to move when stimulated by humans OR collapse when moving OR recumbency OR weight loss >20% of pre-challenge value OR manifestation of any neurologic signs (ataxia, head tilt, etc.) OR body temperature < 100°F; and 5 = found dead. For the challenge, ferrets were lightly anesthetized with ketamine-xylazine, and 0.5 mL containing 10^6^ PFU of influenza A/Vietnam/1203/2004 H5N1 virus was instilled intranasally, split between nares. Following challenge, animals were observed for clinical scores, and temperatures were recorded at least once daily. The animals were weighed at the time of challenge and every day thereafter. Nasal washes were collected under ketamine-xylazine sedation on Days 1, 3, and 5 post-challenge. Lungs were collected after animals succumbed to infection or necropsy. Viral titers were quantified by plaque assay of serial dilution of nasal wash material or processed lung lysate applied to 6-well plates of confluent Madin-Darby canine kidney cells.

### Quantification of IgG and IgA binding titers by ELISA

Quantification of antigen-specific systemic IgG by ELISA was performed on serum as previously described (*32*), and IgA ELISAs were performed on BAL samples. Plates were coated with either H5(H5N1)(A/Vietnam/1203/2004) obtained from Immune Technology (#IT-003-0051p) or H7N9 (A/Anhui/1/2013) HA from Sino Biological (#40103-V08H) followed by blocking for 1 hour. Each serum or BAL sample was diluted 1:4 and then serially 1:2 to create a 14-point dilution curve for each sample. An anti-HA A/Vietnam/1203/04 influenza virus (VN04-8) antibody (Rockland Immunochemicals #200-301-976) was used as a positive control for H5-specific ELISAs. The

Anti-H7N9 Hemagglutinin/HA Antibody (Sino Biological #11082-MM04) was used as a positive control for H7-specific ELISAs. The serially diluted samples were then transferred onto coated, washed plates followed by a 1-hour incubation. For IgG ELISAs, antibodies were detected using the Anti-Mouse IgG (Fc specific)-Alkaline Phosphatase antibody (Sigma-Aldrich #A2429) or an anti-ferret HRP-conjugated antibody (Abcam #ab112770) at a 1:4000 dilution in blocking buffer. For IgA ELISAs, antibodies were detected using the Anti-Mouse IgA (α-chain specific)-Alkaline Phosphatase antibody (Sigma-Aldrich #A4937) at a 1:1000 dilution in blocking buffer. Plates were then developed using the appropriate reactive substrate. Sample concentrations were interpolated off the linear region of each sample dilution curve using the standard curve for absolute quantification of antibody titers, or endpoint titers were calculated by performing a least-squares fit of the OD values at each dilution to a four-parameter sigmoidal curve. LLODs were calculated by taking the slope of the linear portion of the four-point logistic standard curve and dividing that value by three times the standard deviation of the negative control samples.

### Quantification of neutralizing antibodies by pseudovirus neutralization assay

Serum pseudovirus-neutralizing antibodies were quantified as previously described (*32*) with the following modifications. Pseudovirus stocks were generated using HEK-293 cells (American Type Culture Collection #CRL-3216) that were cultured and co-transfected with the following plasmids: a plasmid containing a lentiviral backbone expressing luciferase and ZsGreen (BEI Resources #NR-52516), plasmids containing lentiviral helper genes (BEI Resources #NR-52517, NR-52518, and NR-52519), and a plasmid expressing either H5 A/Vietnam/1203/2004 HA or H7 A/Anhui/2013 HA.

For the pseudovirus neutralization assay, each serum sample was serially diluted and mixed with a viral suspension that had a titer of 5 x 10^7^ fluorescence units/mL. The solution was added to a confluent monolayer of HEK-293T cells plated at 1 x 10^4^ cells/well. Plates were incubated at 37°C and 5% CO_2_ for 3 days. Neutralization was measured via fluorescence in each well. The titers of nAbs in mouse or ferret sera were calculated based on the serum dilution that resulted in half the added virus being neutralized. Pseudovirus neutralization is presented as 50% inhibitory concentration (IC50), which was calculated using a set of virus-infected control wells (100% infectious signal) and media control wells (0% infectious signal), with the limit of detection set based on the reciprocal of the initial serum dilution.

### Statistical analysis

When possible, one-way, parametric statistical comparisons (one-way ANOVA) were performed to assess the statistical difference between groups in the *in vitro* and *in vivo* studies, with a *p* value of 0.05 used as the cut-off. When the above analysis was not possible, such as when the standard deviation was statistically different between groups, a non-parametric analysis was used (Kruskal-Wallis), with a corresponding *p* value of 0.05 as a cutoff. All statistical calculations were performed using GraphPad Prism v10.0. Log-transformed data (antibody titers, viral loads) are displayed with geometric mean and geometric SD, whereas untransformed data (flow cytometric data) show mean and SD.

## ACKNOWLEDGEMENTS

The following reagent was obtained through BEI Resources, NIAID, NIH: A/H5N1 Influenza Vaccine, Inactivated Whole Virion (A/Vietnam/1203/2004), Vero-Cell Derived, Adjuvanted, 15 Micrograms HA, NR-12143. The Influenza Reagent, Influenza Antigen A/Anhui/1/2013 (H7N9) (NIBRG-268), NIBSC code 14/250, was obtained from NIBSC (Potters Bar, Hertfordshire, EN6 3QG). Images for figures were obtained from Clker.com under a CC0 1.0 license (https://creativecommons.org/publicdomain/zero/1.0/) and from Nicolás De Francesco/SciDraw (https://scidraw.io/) and Servier Medical Art (https://smart.servier.com/) under a CC BY 4.0 license (https://creativecommons.org/licenses/by/4.0/). We gratefully acknowledge Dr. Valerie Soza for her support in the preparation of this manuscript for publication and her editorial aid.

## Author contributions

Conceptualization: CC, RAB, EAV

Data curation: MRY, DNK

Formal analysis: MRY, MAD, DNK, MFJ

Funding acquisition: CC, AG, RAB, EAV

Investigation: MRY, DNK, MFJ, EL, JS, SB, NC, EM, SR, CP, PF, JB, ATH

Methodology: MRY, DNK, EL, SB, NC

Project administration: MAD, AG, RAB, EAV

Resources: CC, AG, RAB, EAV

Supervision: EM, AG, RAB, EAV

Validation: MRY, DNK, MFJ

Visualization: MRY, DNK

Writing – original draft: MRY, MAD

Writing – review & editing: MRY, MAD, DNK, MFJ, EL, JS, SB, NC, EM, SR, CP, PF, JB, CC, ATH, AG, RAB, EAV

## Data and materials availability

The datasets generated and/or analyzed during the current study are available from the corresponding author on reasonable request. AAHI’s unique saRNA constructs and novel Nanostructured Lipid Carrier (NLC) are available for noncommercial research or scholarly purposes under the terms and conditions of an appropriate material transfer agreement. Correspondence and material requests should be addressed to the corresponding author.

## SUPPLEMENTARY MATERIALS

Supplementary Figure S1. Biophysical characterization and additional immune readouts for monovalent H5 saRNA-NLC vaccine.

Supplementary Figure S2. Additional data from H5 saRNA-NLC vaccine monovalent dosing study relevant to Figure 1.

Supplementary Figure S3. Biophysical characterization and additional immune readouts for monovalent H7 saRNA-NLC vaccine.

Supplementary Figure S4. Additional data from H7 saRNA-NLC vaccine monovalent dosing study relevant to Figure 2.

Supplementary Figure S5. Biophysical characterization and additional immune readouts for bivalent H5/H7 saRNA-NLC vaccine.

Supplementary Figure S6. Full bivalent intramuscular study systemic immunogenicity data relevant to Figure 3.

Supplementary Figure S7. Full bivalent intramuscular study mucosal immunogenicity data relevant to Figure 3.

Supplementary Figure S8. Full bivalent intranasal study systemic immunogenicity data relevant to Figure 4.

Supplementary Figure S9. Full bivalent intranasal study mucosal immunogenicity data relevant to Figure 4.

Supplementary Figure S10. Additional systemic immunogenicity data for the routes of bivalent vaccination study presented in Figure 5.

Supplementary Figure S11. Additional mucosal immunogenicity data for the routes of bivalent vaccination study presented in Figure 5.

Supplementary Figure S12. Ferret body temperature after H7N9 challenge.

Supplementary Figure S13. Additional metrics for tracking ferret outcomes post-H5N1 challenge.

Supplementary Figure S14. Representative splenic T cell ICS flow cytometry gating strategy.

Supplementary Figure S15. Representative lung T cell ICS flow cytometry gating strategy.

