## Supplementary Materials for "An intranasal, NLC-delivered self-amplifying RNA vaccine establishes protective immunity against pre-pandemic H5N1 and H7N9 influenza"

### Supplementary Figures

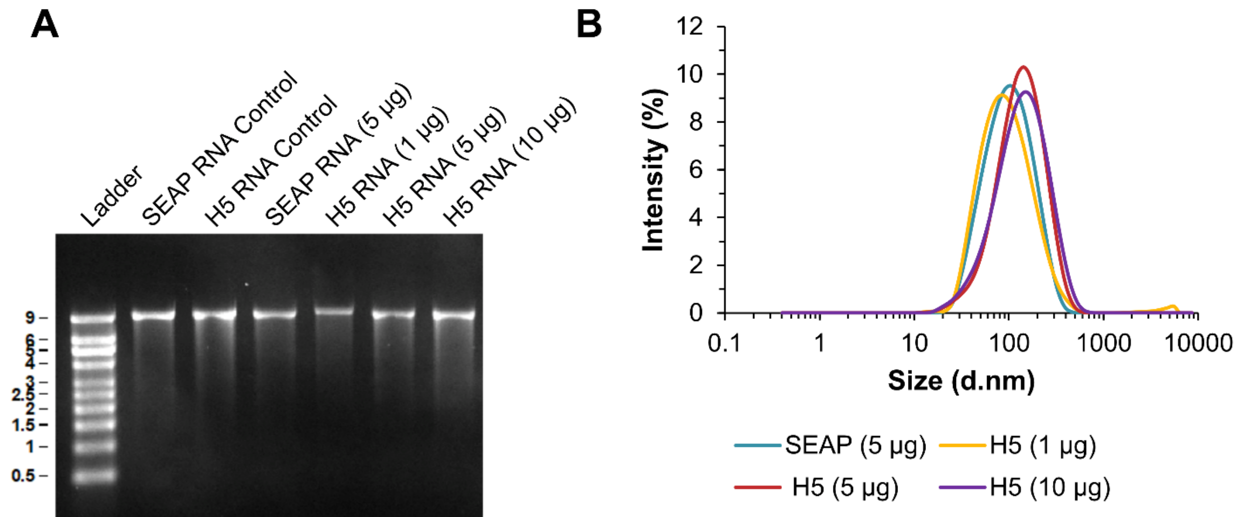


| RNA | Z-Average (d.nm) | PdI |
|---|---|---|
| SEAP (5 µg) | 85 ± 1.1 | 0.23 ± 0.01 |
| H5 (1 µg) | 81 ± 3.3 | 0.25 ± 0.02 |
| H5 (5 µg) | 113 ± 3.4 | 0.24 ± 0.01 |
| H5 (10 µg) | 112 ± 3.4 | 0.27 ± 0.00 |

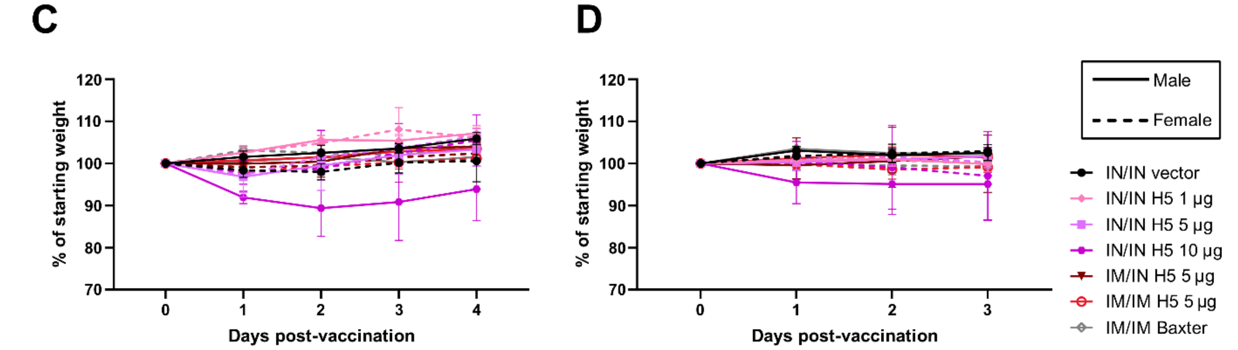


**Supplementary Figure S1. Biophysical characterization and additional immune readouts for monovalent H5 saRNA-NLC vaccine.** (**A**) RNA size and integrity before complexing ("Control") and after extraction from the saRNA-NLC complexes as measured by agarose gel electrophoresis. (**B**) Average size and polydispersity index (PdI) for saRNA-NLC vaccine complexes, after measurement in triplicate. Particle size intensity distribution of saRNA-NLC vaccine complexes as measured by dynamic light scattering. (**C-D**) Mouse body weights (**C**) post-prime and (**D**) post-boost. *n* = 6 (3M:3F) for vector control (SEAP) group and *n* = 8 (4M:4F) for experimental groups.

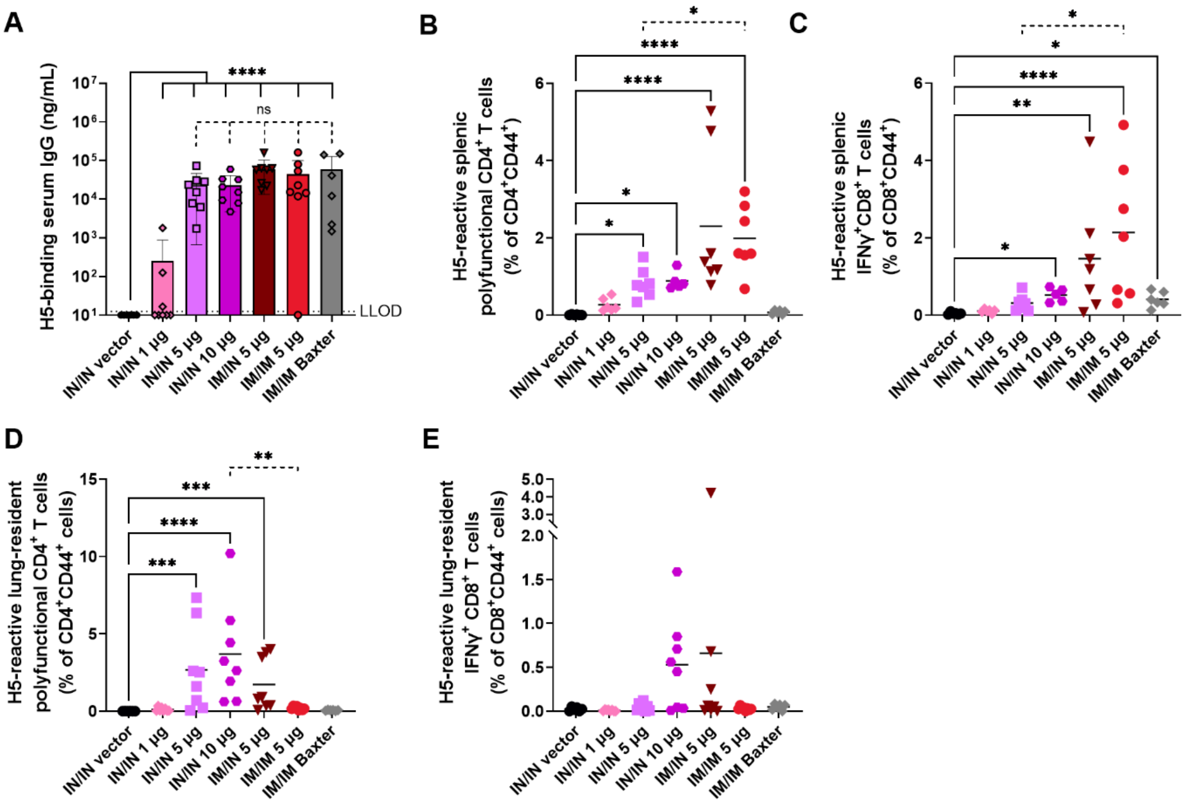


**Supplementary Figure S2. Additional data from H5 saRNA-NLC vaccine monovalent dosing study relevant to Figure 1.** (A) Post-prime serum H5-binding IgG titers. Post-boost H5-reactive (B) splenic polyfunctional (IFN $\gamma^+$ IL-$2^+$ TNF $\alpha^+$) $CD4^+$ T cells, (C) splenic IFN $\gamma^+$ $CD8^+$ T cells, (D) lung-resident ($CD69^+$) polyfunctional (IFN $\gamma^+$ IL-$2^+$ TNF $\alpha^+$) $CD4^+$ T cells, and (E) lung-resident ($CD69^+$ $CD103^+$) IFN $\gamma^+$ $CD8^+$ T cells. (B-E) For all assays, $n$ = 6 for vector control group and $n$ = 8 for experimental groups. All groups were sex balanced. Two statistical hypotheses were tested within each figure, with filled lines showing comparisons to the vector control and dotted lines representing tests between key experimental groups. (A) Statistical analyses performed on log-transformed data using one-way ANOVA with Šídák's multiple comparison test. (B-E) Statistical analysis represents Kruskal-Wallis test with Dunn's multiple comparisons (filled lines and dotted lines). LLOD = lower limit of detection; ns = not significant. * $p < 0.05$, ** $p < 0.01$, *** $p < 0.001$, **** $p < 0.0001$.

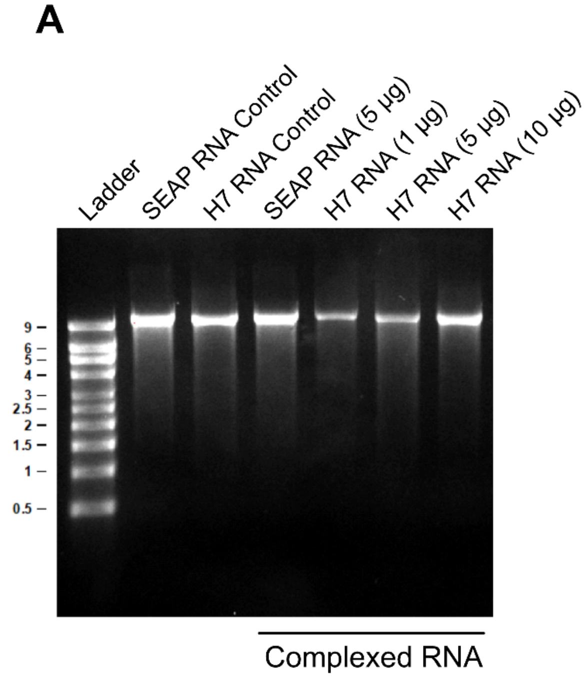


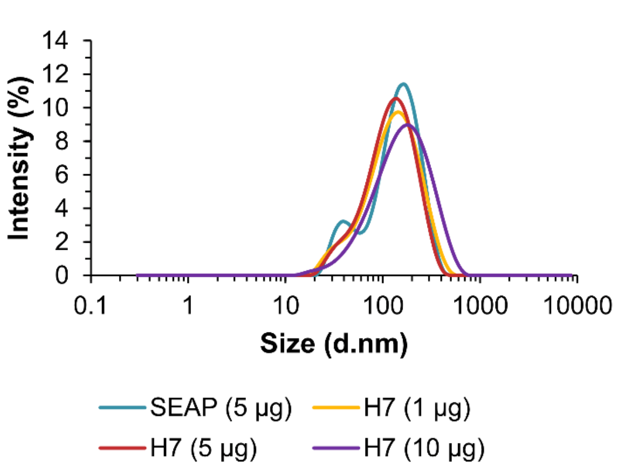


| RNA | Z-Average (d.nm) | PdI |
|---|---|---|
| SEAP (5 µg) | 107 ± 1.9 | 0.27 ± 0.01 |
| H7 (1 µg) | 104 ± 0.5 | 0.27 ± 0.01 |
| H7 (5 µg) | 100 ± 1.5 | 0.25 ± 0.01 |
| H7 (10 µg) | 127 ± 2.7 | 0.28 ± 0.01 |

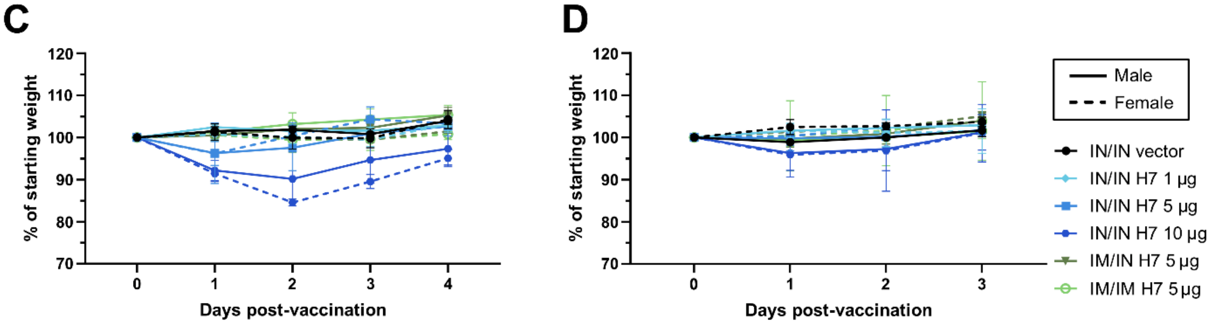


**Supplementary Figure S3. Biophysical characterization and additional immune readouts for monovalent H7 saRNA-NLC vaccine.** (**A**) RNA size and integrity before complexing ("Control") and after extraction from the saRNA-NLC complexes as measured by agarose gel electrophoresis. (**B**) Average size and polydispersity index (PdI) for saRNA-NLC vaccine complexes, after measurement in triplicate. Particle size intensity distribution of saRNA-NLC vaccine complexes as measured by dynamic light scattering. (**C-D**) Mouse body weights (**C**) post-prime and (**D**) post-boost. *n* = 6 (3M:3F) for vector control (SEAP) group and *n* = 8 (4M:4F) for experimental groups.

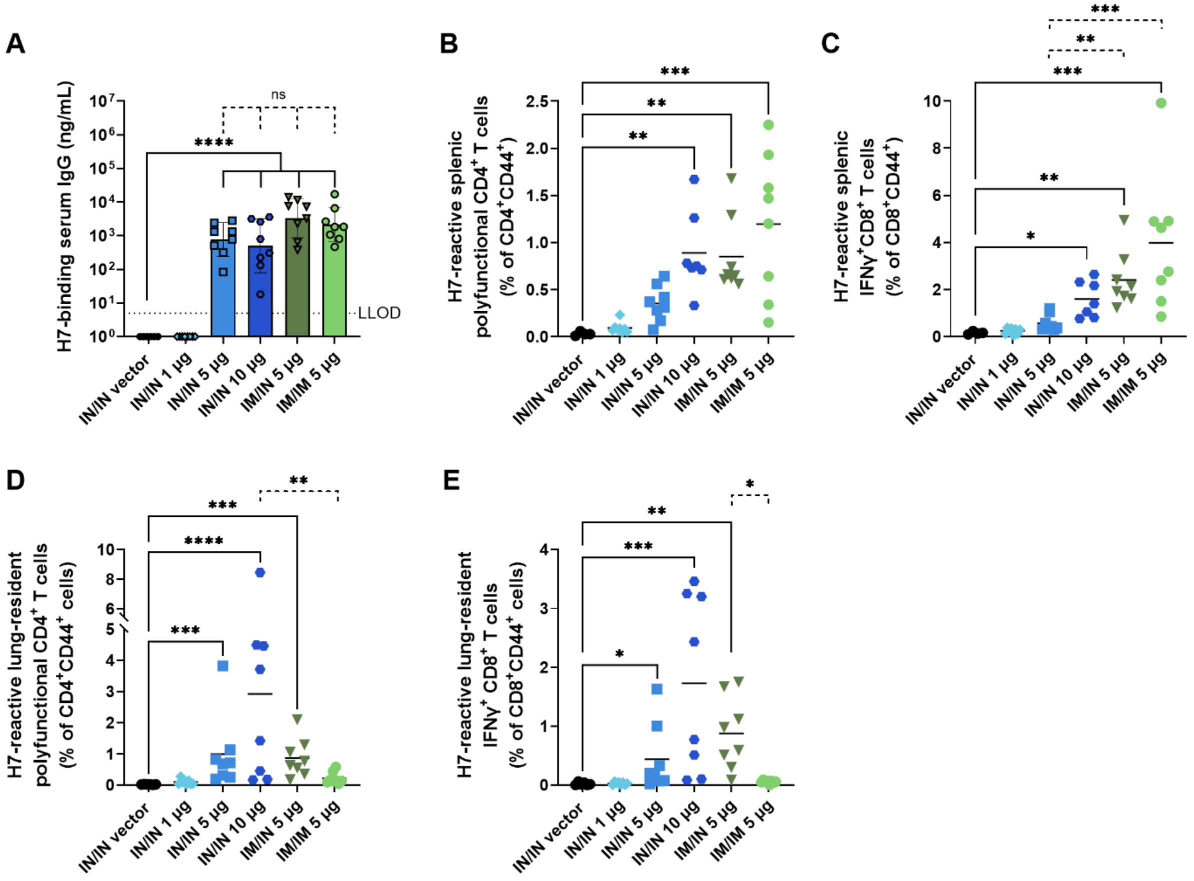


**Supplementary Figure S4. Additional data from H7 saRNA-NLC vaccine monovalent dosing study relevant to Figure 2.** (A) Post-prime serum H7-binding IgG titers. Post-boost H7-reactive (B) splenic polyfunctional (IFN$\gamma^+$ IL-$2^+$ TNF$\alpha^+$) CD$4^+$ T cells, (C) splenic IFN$\gamma^+$ CD$8^+$ T cells, (D) lung-resident (CD$69^+$) polyfunctional (IFN$\gamma^+$ IL-$2^+$ TNF$\alpha^+$) CD$4^+$ T cells, and (E) lung-resident (CD$69^+$ CD$103^+$) IFN$\gamma^+$ CD$8^+$ T cells. For all assays, $n = 6$ for vector control group, $n = 8$ for experimental groups. All groups were sex balanced. Two statistical hypotheses were tested within each figure, with filled lines showing comparisons to the vector control and dotted lines representing tests between key experimental groups. (A) Statistical analyses performed on log-transformed data using one-way ANOVA with Šídák's multiple comparison test. (B-E) Statistical analysis represents Kruskal-Wallis test with Dunn's multiple comparisons (filled lines [B-E] and dotted lines [C, D]) or Brown-Forsythe and Welch ANOVA with Dunnett's T3 multiple comparisons (dotted lines [B, E]). LLOD = lower limit of detection; ns = not significant. * $p < 0.05$, ** $p < 0.01$, *** $p < 0.001$, **** $p < 0.0001$.

**A**

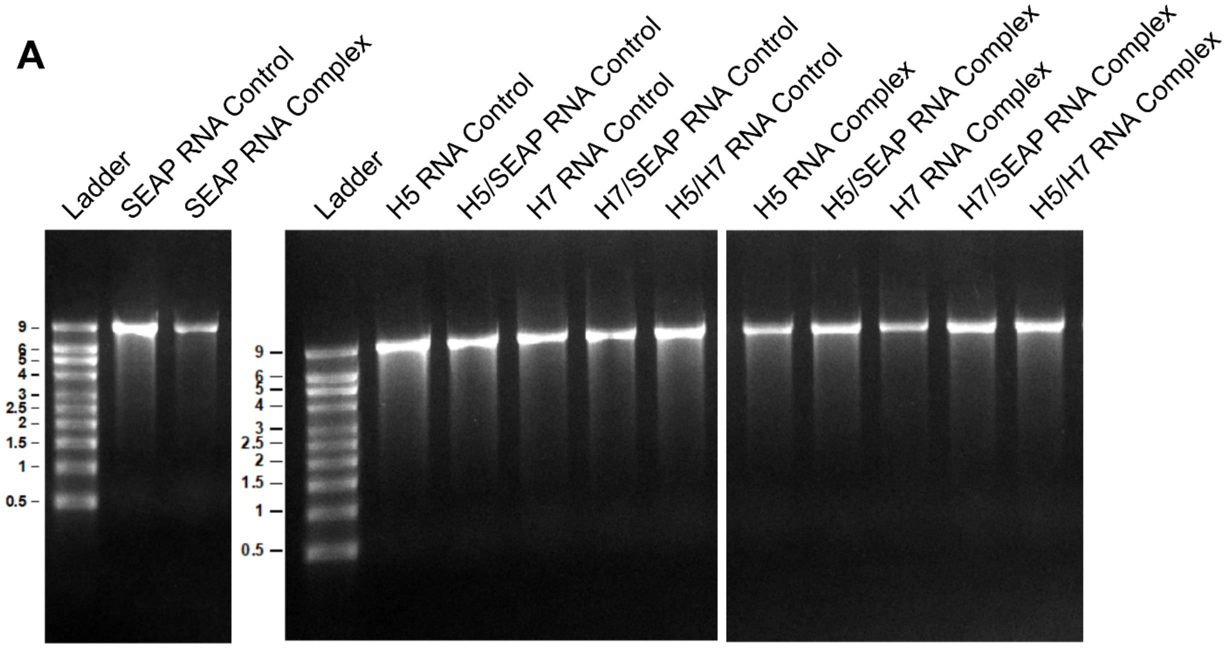


**B**

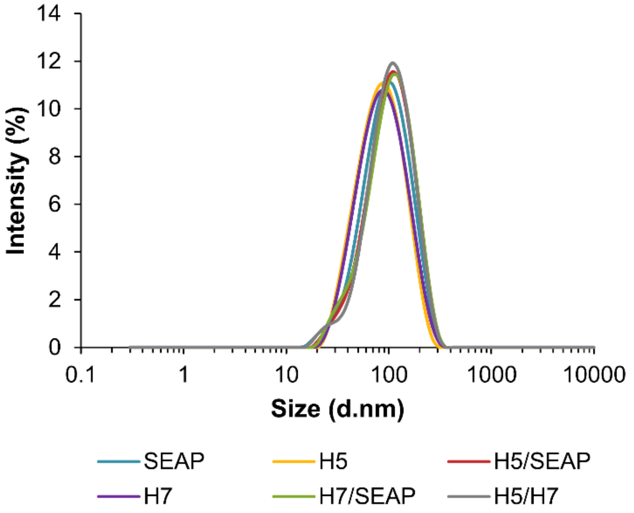


| RNA | Z-Average (d.nm) | PdI |
|---|---|---|
| SEAP | 90± 1.7 | 0.24± 0.01 |
| H5 | 85± 2.0 | 0.23± 0.01 |
| H5/SEAP | 80± 1.4 | 0.22± 0.00 |
| H7 | 82± 1.2 | 0.22± 0.01 |
| H7/SEAP | 74± 1.9 | 0.19± 0.01 |
| H5/H7 | 88± 1.6 | 0.21± 0.01 |

**Supplementary Figure S5. Biophysical characterization and additional immune readouts for bivalent H5/H7 saRNA-NLC vaccine.** (**A**) RNA size and integrity before complexing ("Control") and after extraction from the saRNA-NLC complex ("Complex") as measured by agarose gel electrophoresis. (**B**) Average size and polydispersity index (PdI) for saRNA-NLC vaccine complexes, after measurement in triplicate. Particle size intensity distribution of saRNA-NLC vaccine complexes as measured by dynamic light scattering. Bivalent vaccines (H5/SEAP, H7/SEAP, and H5/H7) each represent 10 µg saRNA total (5 µg of each saRNA-NLC vaccine combined by simple mixing).

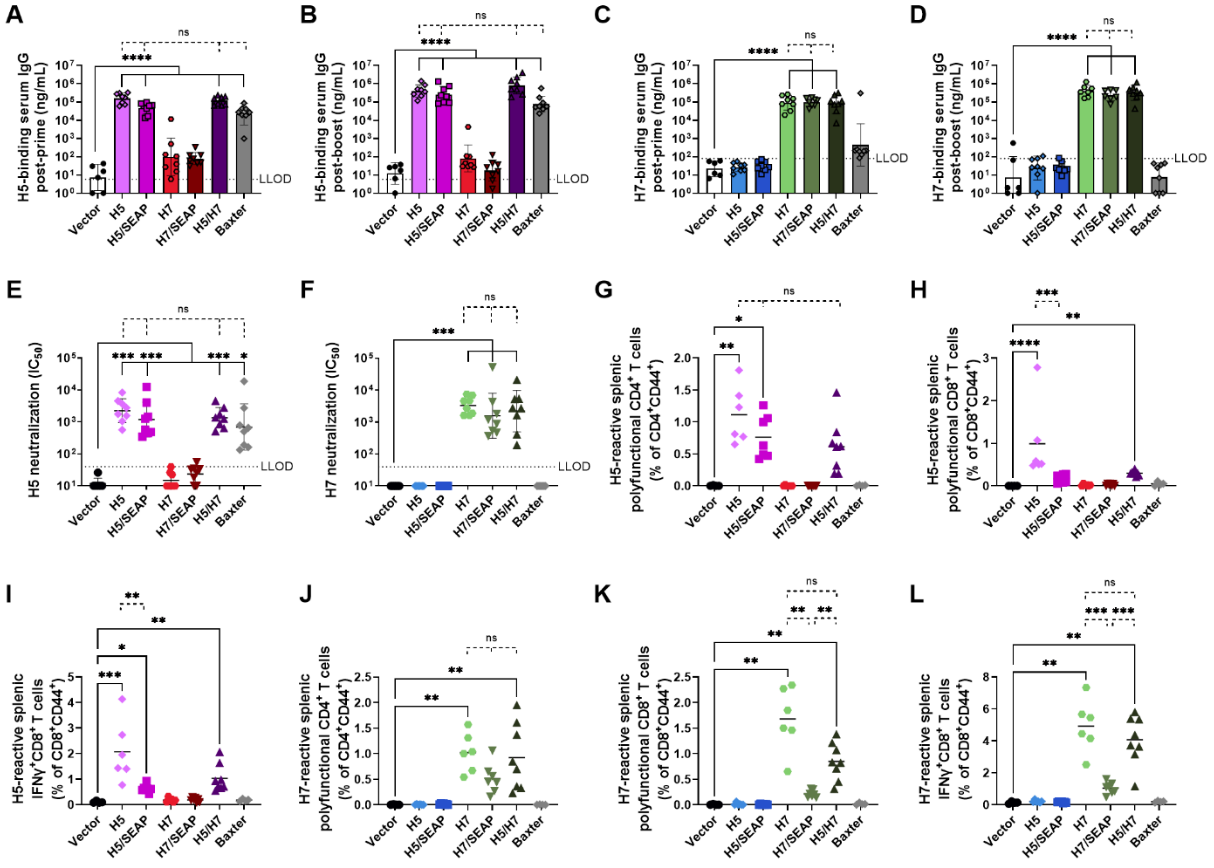


**Supplementary Figure S6. Full bivalent intramuscular study systemic immunogenicity data relevant to Figure 3.** (A-B) H5 and (C-D) H7 HA-binding IgG (A, C) post-prime and (B, D) post-boost serum titers. (E) H5N1 and (F) H7N9 neutralization capacity ($IC_{50}$) of post-boost mouse serum. (G-I) Post-boost H5-reactive splenic (G) polyfunctional (IFN$\gamma^{+}$ IL-2$^{+}$ TNF$\alpha^{+}$) CD4$^{+}$ T cells, (H) polyfunctional (IFN$\gamma^{+}$ IL-2$^{+}$ TNF$\alpha^{+}$) CD8$^{+}$ T cells, and (I) IFN$\gamma^{+}$ CD8$^{+}$ T cells. (J-L) Post-boost H7-reactive splenic (J) polyfunctional (IFN$\gamma^{+}$ IL-2$^{+}$ TNF$\alpha^{+}$) CD4$^{+}$ T cells, (K) polyfunctional (IFN$\gamma^{+}$ IL-2$^{+}$ TNF$\alpha^{+}$) CD8$^{+}$ T cells, and (L) IFN$\gamma^{+}$ CD8$^{+}$ T cells. For all assays, $n = 6$ animals for vector control (SEAP) group, $n = 8$ for experimental groups. All groups were sex balanced. Two statistical hypotheses were tested within each figure, unless specifically listed, with filled lines showing comparisons to the vector control and dotted lines representing tests between key experimental groups. (A-D) Statistical analyses performed on log-transformed data used one-way ANOVA with Šídák's multiple comparison test. (E-F) Statistical analysis performed on log-transformed data used Kruskal-Wallis test with Dunn's multiple comparisons. (G-J) Statistical analysis represents Kruskal-Wallis test with Dunn's multiple comparisons (filled lines [G-J] and dotted lines [H, I]), one-way ANOVA with Tukey's multiple comparisons (dotted lines [G, J, L]), or Brown-Forsythe and Welch ANOVA with Dunnett's T3 multiple comparisons (dotted lines [K]). -LLOD = lower limit of detection; ns = not significant. * $p < 0.05$, ** $p < 0.01$, *** $p < 0.001$, **** $p < 0.0001$.

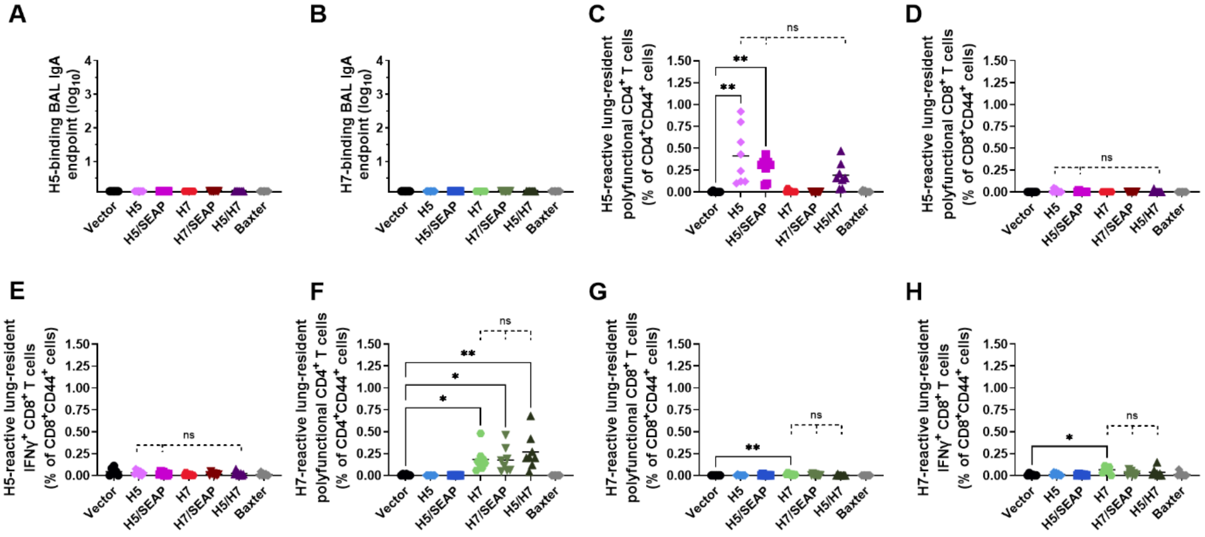


**Supplementary Figure S7. Full bivalent intramuscular study mucosal immunogenicity data relevant to Figure 3.** Post-boost (A) H5 and (B) H7 HA-binding IgA antibody titers in BAL samples. (C-E) Post-boost H5-reactive lung-resident ($CD69^+$) (C) polyfunctional ($IFN\gamma^+$ $IL\text{-}2^+$ $TNF\alpha^+$) $CD4^+$ T cells, lung-resident ($CD69^+$ $CD103^+$) (D) polyfunctional ($IFN\gamma^+$ $IL\text{-}2^+$ $TNF\alpha^+$) $CD8^+$ T cells, and lung-resident ($CD69^+$ $CD103^+$) (E) $IFN\gamma^+$ $CD8^+$ T cells. (F-H) Post-boost H7-reactive lung-resident ($CD69^+$) (F) polyfunctional ($IFN\gamma^+$ $IL\text{-}2^+$ $TNF\alpha^+$) $CD4^+$ T cells, lung-resident ($CD69^+$ $CD103^+$) (G) polyfunctional ($IFN\gamma^+$ $IL\text{-}2^+$ $TNF\alpha^+$) $CD8^+$ T cells, and lung-resident ($CD69^+$ $CD103^+$) (H) $IFN\gamma^+$ $CD8^+$ T cells. For all assays, $n = 6$ animals for vector control (SEAP) group, $n = 8$ for experimental groups. All groups were sex balanced. Two statistical hypotheses were tested within each figure, unless specifically listed, with filled lines showing comparisons to the vector control) and dotted lines representing tests between key experimental groups. (C-H) Statistical analysis represents Kruskal-Wallis test with Dunn's multiple comparisons (filled lines [C-H] and dotted lines [D, F-H]), one-way ANOVA with Tukey's multiple comparisons (dotted lines [E]), or Brown-Forsythe and Welch ANOVA with Dunnett's T3 multiple comparisons (dotted lines [C]). ns = not significant. * $p < 0.05$, ** $p < 0.01$.

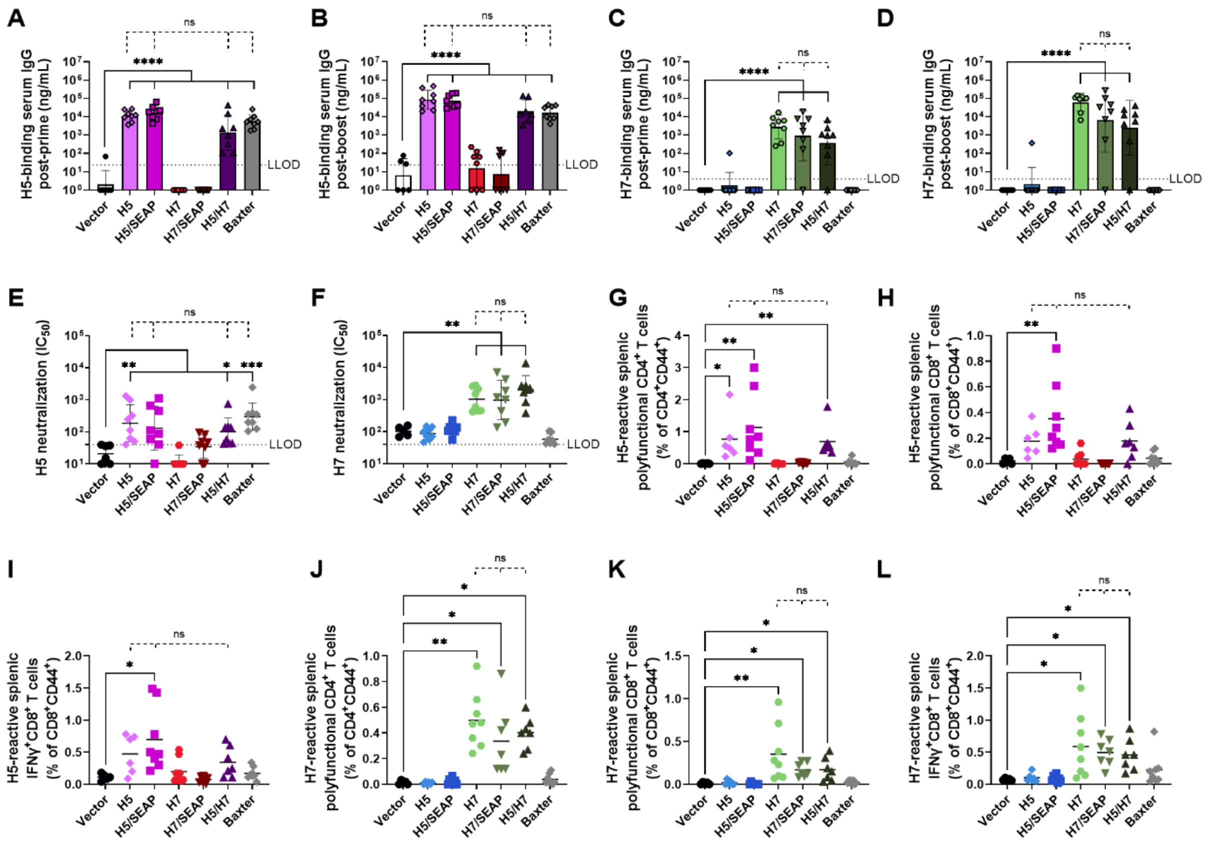

**Supplementary Figure S8. Full bivalent intranasal study systemic immunogenicity data relevant to Figure 4.** Serum (A-B) H5 and (C-D) H7-binding IgG titers (A, C) post-prime and (B, D) post-boost. (E) Post-boost serum (E) H5 and (F) H7 pseudovirus neutralization capacity ($IC_{50}$). (G-I) Post-boost H5-reactive splenic (G) polyfunctional ($IFN\gamma^{+}$ $IL\text{-}2^{+}$ $TNF\alpha^{+}$) $CD4^{+}$ T cells, (H) polyfunctional ($IFN\gamma^{+}$ $IL\text{-}2^{+}$ $TNF\alpha^{+}$) $CD8^{+}$T cells, and (I) $IFN\gamma^{+}$ $CD8^{+}$ T cells. (J-L) Post-boost H7-reactive splenic (J) polyfunctional ($IFN\gamma^{+}$ $IL\text{-}2^{+}$ $TNF\alpha^{+}$) $CD4^{+}$ T cells, (K) polyfunctional ($IFN\gamma^{+}$ $IL\text{-}2^{+}$ $TNF\alpha^{+}$) $CD8^{+}$ T cells, and (L) $IFN\gamma^{+}$ $CD8^{+}$ T cells. For all assays, $n = 6$ animals for vector control (SEAP) group, $n = 8$ for experimental groups. All groups were sex balanced. Two statistical hypotheses were tested within each figure, unless specifically listed, with filled lines showing comparisons to the vector control and dotted lines representing tests between key experimental groups. (A-D) Statistical analyses performed on log-transformed data used one-way ANOVA with Šídák's multiple comparison test. (E-F) Statistical analysis performed on log-transformed data used Kruskal-Wallis test with Dunn's multiple comparison test. (G-L) Statistical analysis represents Kruskal-Wallis test with Dunn's multiple comparisons (filled lines [G-L] and dotted lines [G, H]) or one-way ANOVA with Tukey's multiple comparisons (dotted lines [I-L]). LLOD = lower limit of detection; ns = not significant. * $p < 0.05$, ** $p < 0.01$, *** $p < 0.001$, **** $p < 0.0001$.

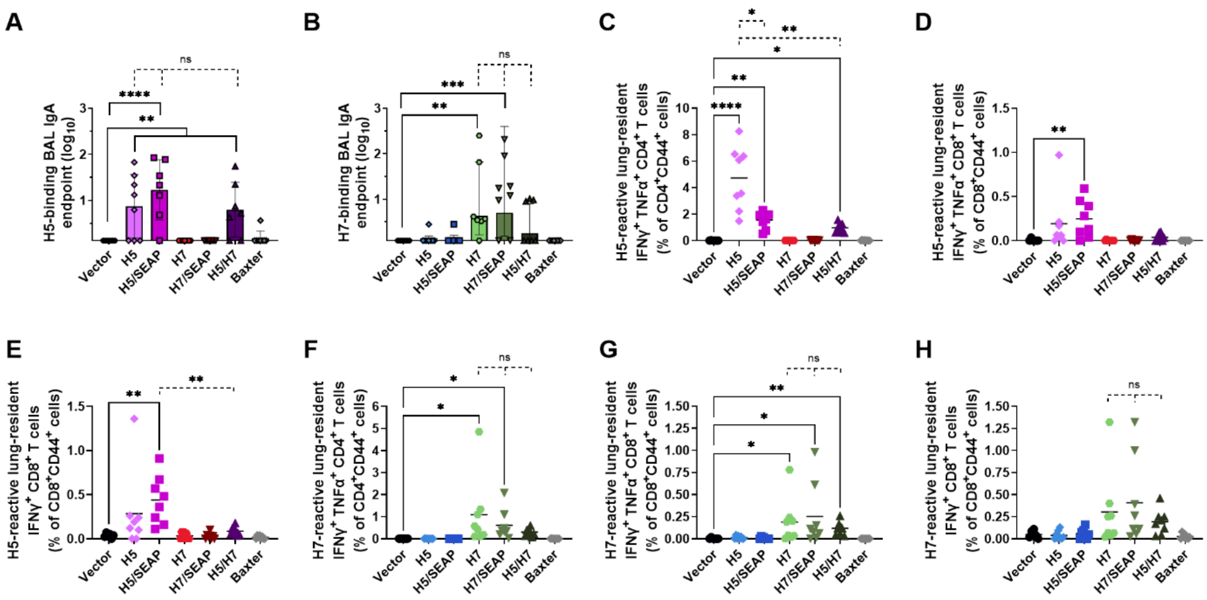

**Supplementary Figure S9. Full bivalent intranasal study mucosal immunogenicity data relevant to Figure 4.** Post-boost (A) H5 and (B) H7 HA-binding IgA antibody titers in BAL samples. (C-E) Post-boost H5-reactive lung-resident ($CD69^+$) (C) bifunctional ($IFN\gamma^+$ $TNF\alpha^+$) $CD4^+$ T cells, lung-resident ($CD69^+$ $CD103^+$) (D) bifunctional ($IFN\gamma^+$ $TNF\alpha^+$) $CD8^+$ T cells, and lung-resident ($CD69^+$ $CD103^+$) (E) $IFN\gamma^+$ $CD8^+$ T cells. (F-H) Post-boost H7-reactive lung-resident ($CD69^+$) (F) bifunctional ($IFN\gamma^+$ $TNF\alpha^+$) $CD4^+$ T cells, lung-resident ($CD69^+$ $CD103^+$) (G) bifunctional ($IFN\gamma^+$ $TNF\alpha^+$) $CD8^+$ T cells, and lung-resident ($CD69^+$ $CD103^+$) (H) $IFN\gamma^+$ $CD8^+$ T cells. For all assays, $n = 6$ animals for vector control (SEAP) group, $n = 8$ for experimental groups. All groups were sex balanced. Two statistical hypotheses were tested within each figure, with filled lines showing comparisons to the vector control and dotted lines representing tests between key experimental groups. (A-B) Statistical analyses performed on log-transformed data used one-way ANOVA with Šídák's multiple comparison test. (C-H) Statistical analysis represents Kruskal-Wallis test with Dunn's multiple comparisons (filled lines [C-H] and dotted lines [D-H]) or Brown-Forsythe and Welch ANOVA with Dunnett's T3 multiple comparisons (dotted lines [C]). ns = not significant. * $p < 0.05$, ** $p < 0.01$, *** $p < 0.001$, **** $p < 0.0001$.

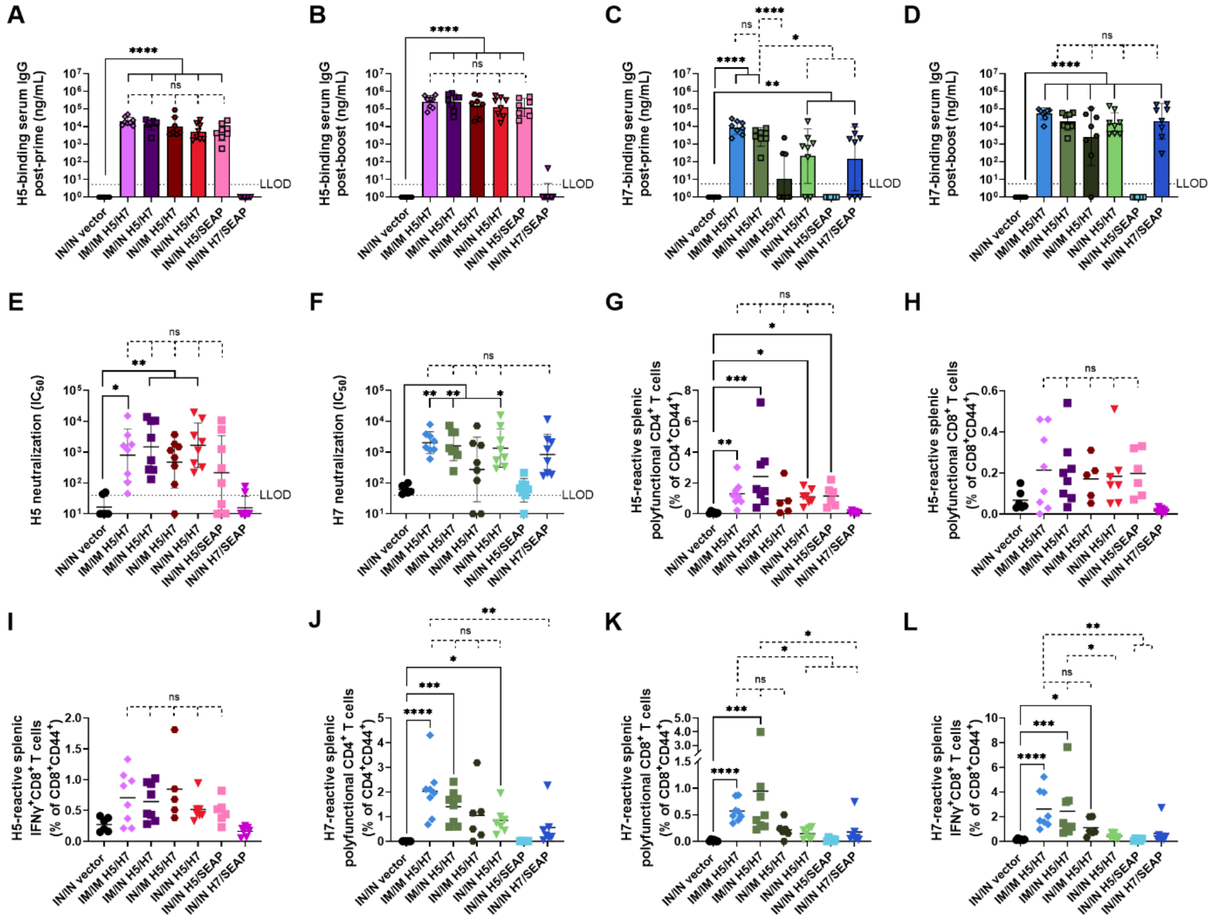


**Supplementary Figure S10. Additional systemic immunogenicity data for the routes of bivalent vaccination study presented in Figure 5.** (A-B) H5 and (C-D) H7 HA-binding IgG (A, C) post-prime and (B, D) post-boost serum IgG titers. (E) H5N1 and (F) H7N9 neutralization capacity ($IC_{50}$) of post-boost mouse serum. (G-I) Post-boost H5-reactive splenic (G) polyfunctional ($IFN\gamma^{+}$ $IL\text{-}2^{+}$ $TNF\alpha^{+}$) $CD4^{+}$ T cells, (H) polyfunctional ($IFN\gamma^{+}$ $IL\text{-}2^{+}$ $TNF\alpha^{+}$) $CD8^{+}$ T cells, and (I) $IFN\gamma^{+}$ $CD8^{+}$ T cells. (J-L) Post-boost H7-reactive splenic (J) polyfunctional ($IFN\gamma^{+}$ $IL\text{-}2^{+}$ $TNF\alpha^{+}$) $CD4^{+}$ T cells, (K) polyfunctional ($IFN\gamma^{+}$ $IL\text{-}2^{+}$ $TNF\alpha^{+}$) $CD8^{+}$ T cells, and (L) $IFN\gamma^{+}$ $CD8^{+}$ T cells. For all assays, $n = 6$ animals for vector control (SEAP) group, $n = 8$ for experimental groups. All groups were sex balanced. Two statistical hypotheses were tested within each figure, unless specifically listed, with filled lines showing comparisons to the vector control and dotted lines representing tests between key experimental groups. (A-D) Statistical analyses performed on log-transformed data used one-way ANOVA with Šídák's multiple comparison test. (E-F) Statistical analysis performed on log-transformed data used Kruskal-Wallis test with Dunn's multiple comparisons. (G-L) Statistical analysis represents Kruskal-Wallis test with Dunn's multiple comparisons (filled lines [G, H, J-L] and dotted lines [G, J-L]), one-way ANOVA with Tukey's multiple comparisons (dotted lines [H, I]), or Brown-Forsythe and Welch ANOVA with Dunnett's T3 multiple comparisons (filled lines [I]). LLOD = lower limit of detection; ns = not significant. * $p < 0.05$, ** $p < 0.01$, *** $p < 0.001$, **** $p < 0.0001$.

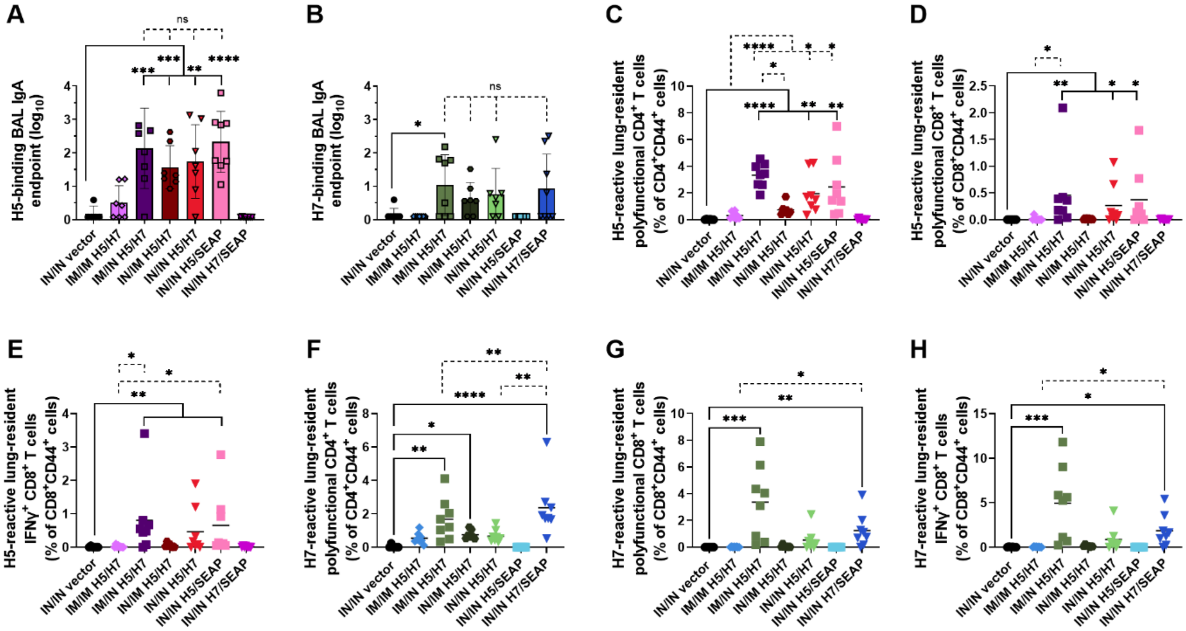


**Supplementary Figure S11. Additional mucosal immunogenicity data for the routes of bivalent vaccination study presented in Figure 5.** (A) H5 and (B) H7 HA-binding IgA antibody titers in mouse BAL material post-boost. (C-E) Post-boost H5-reactive lung-resident ($CD69^+$) (C) polyfunctional ($IFN\gamma^+$ $IL\text{-}2^+$ $TNF\alpha^+$) $CD4^+$ T cells, lung-resident ($CD69^+$ $CD103^+$) (D) polyfunctional ($IFN\gamma^+$ $IL\text{-}2^+$ $TNF\alpha^+$) $CD8^+$ T cells, and lung-resident ($CD69^+$ $CD103^+$) (E) $IFN\gamma^+$ $CD8^+$ T cells. (F-H) Post-boost H7-reactive lung-resident ($CD69^+$) (F) polyfunctional ($IFN\gamma^+$ $IL\text{-}2^+$ $TNF\alpha^+$) $CD4^+$ T cells, lung-resident ($CD69^+$ $CD103^+$) (G) polyfunctional ($IFN\gamma^+$ $IL\text{-}2^+$ $TNF\alpha^+$) $CD8^+$ T cells, and lung-resident ($CD69^+$ $CD103^+$) (H) $IFN\gamma^+$ $CD8^+$ T cells. For all assays, $n = 6$ animals for vector control (SEAP) group, $n = 8$ for experimental groups. All groups were sex balanced. Two statistical hypotheses were tested within each figure, with filled lines showing comparisons to the vector control and dotted lines representing tests between key experimental groups. (A-B) Statistical analyses performed on log-transformed data used one-way ANOVA with Šídák's multiple comparison test. (C-H) Statistical analysis represents Kruskal-Wallis test with Dunn's multiple comparisons (filled lines and dotted lines). ns = not significant. * $p < 0.05$, ** $p < 0.01$, *** $p < 0.001$, **** $p < 0.0001$.

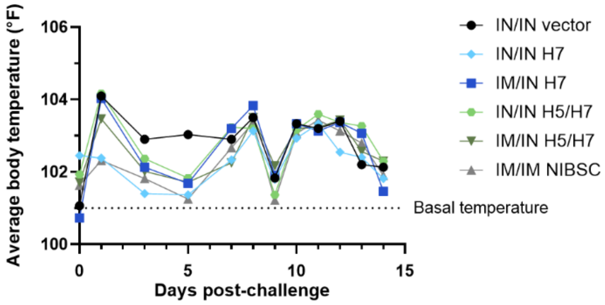


**Supplementary Figure S12. Ferret body temperature after H7N9 challenge**. *n* = 6 (3 male and 3 female) animals per group. NIBSC = NIBSC Influenza Antigen A/Anhui/1/2013 (H7N9).

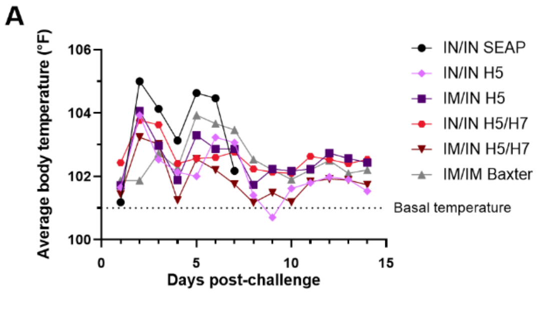


**Supplementary Figure S13. Additional metrics for tracking ferret outcomes post-H5N1 challenge. (A)** Ferret body temperature. $n = 6$ (3 male and 3 female) animals per group. **(B-C)** Viral titers in lung tissue at necropsy.

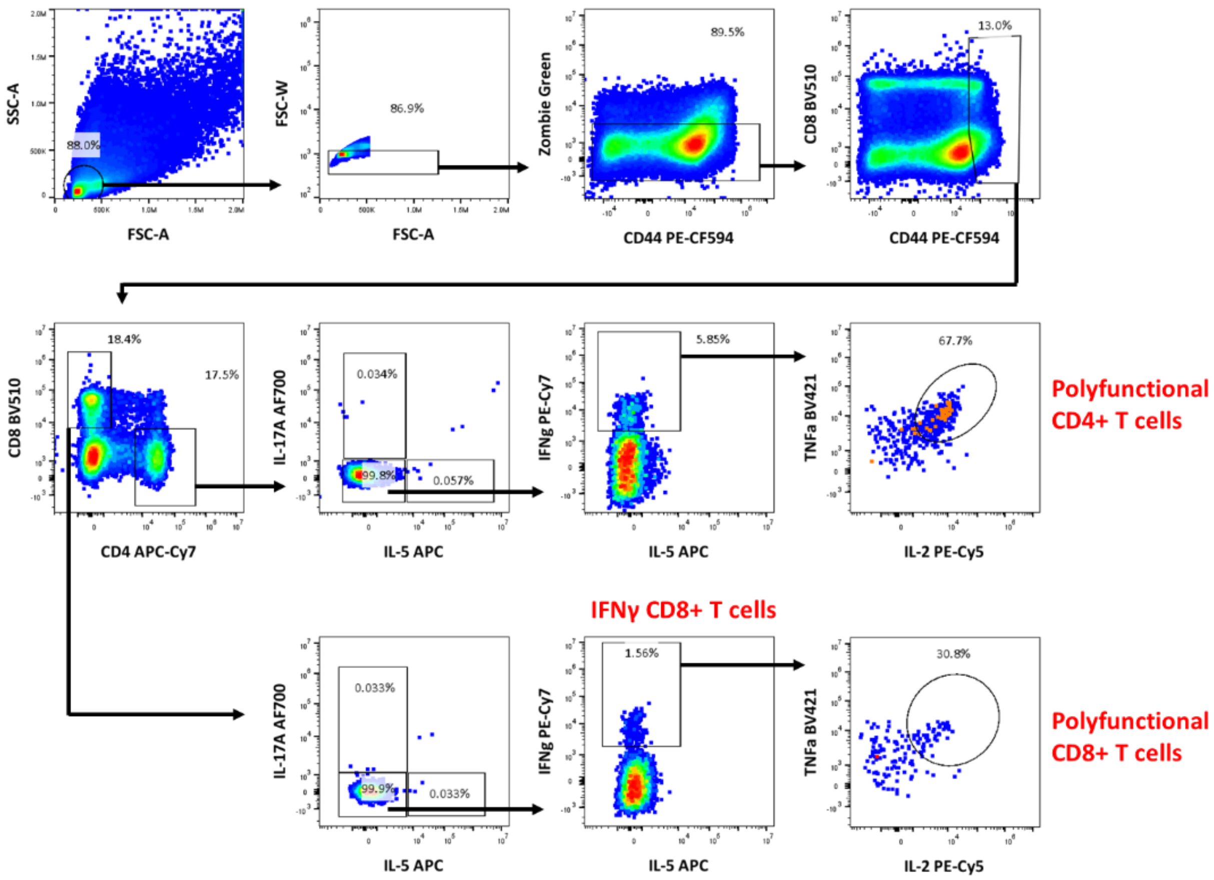


**Supplementary Figure S14. Representative splenic T cell ICS flow cytometry gating strategy.** Biplots depict the gating strategy used to identify polyfunctional ($TNF\alpha^+$ $IL\text{-}2^+$ $IFN\gamma^+$) $CD4^+$ T cells, polyfunctional $CD8^+$ T cells, and $IFN\gamma^+$ $CD8^+$ T cells isolated from mouse spleens. Data shown are from splenocytes stimulated *in vitro* with an H7 peptide pool following prime/boost IM/IM immunization with the H5/H7 bivalent saRNA-NLC vaccine.

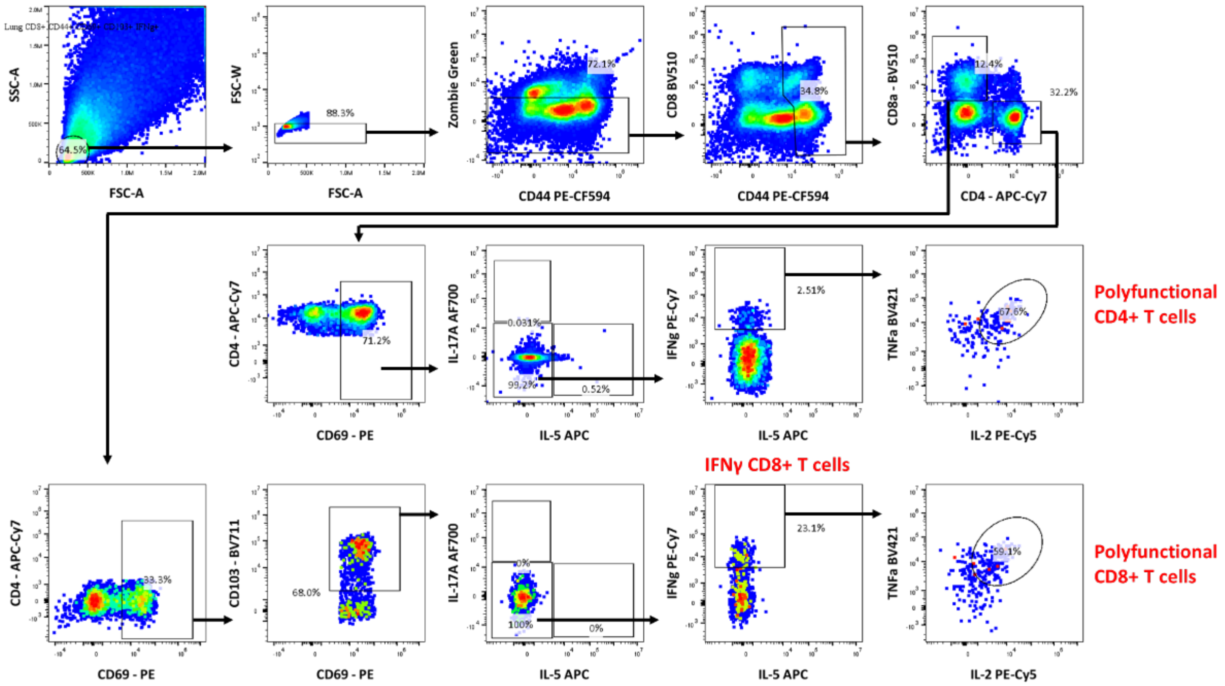


**Supplementary Figure S15. Representative lung T cell ICS flow cytometry gating strategy.** Biplots depict the gating strategy used to identify polyfunctional (TNFα$^+$ IL-2$^+$ IFNγ$^+$) CD4$^+$ T cells, polyfunctional CD8$^+$ T cells, and IFNγ$^+$ CD8$^+$ T cells isolated from mouse lungs. Data shown are from lung lymphocytes stimulated *in vitro* with an H7 peptide pool following prime/boost IM/IN immunization with the H5/H7 bivalent saRNA-NLC vaccine.
